# Proteome-wide profiling reveals compartment-specific RNA buffering of protein aggregation during stress

**DOI:** 10.64898/2026.11.04.764228

**Authors:** Johanna Franziska Seidler, Fabienne Thelen, Nadine Bianca Wäber, Jan Weber, Leon Lohle, Thomas Timm, Günter Lochnit, Jochen Blom, Alexander Goesmann, Katja Sträßer, Cornelia Kilchert

**Author notes:** The first two authors should be regarded as Joint First Authors.

## Abstract

Cellular adaptation to stress requires extensive remodeling of RNA metabolism. The rewiring of the RNA-protein interaction landscape and the assembly of biomolecular condensates are central to this response, yet the global stress-induced reorganization of macromolecular RNA-protein assemblies remains incompletely characterized. Here, we apply R-DeeP, a density gradient approach coupled to quantitative mass spectrometry, to profile the RNA-dependent sedimentation behaviour of the *Saccharomyces cerevisiae* proteome during heat and salt stress. We demonstrate that stress triggers a profound, stress-specific reorganization of the RNA interactome. Many proteins undergo pronounced, selective shifts into high-molecular weight fractions, indicative of aggregation. RNase treatment markedly exacerbates this aggregation, consistent with the model that RNA acts as a general chaperone that keeps proteins with intrinsically disordered regions soluble. Strikingly, our data demonstrate that this protection is compartmentalised. Nuclear proteins are partially shielded from stress-induced aggregation. This protection coincides with the nuclear accumulation of RNA caused by a block of RNA export during stress, and RNase treatment abolishes this protection. Nuclear RNA retention may therefore act as a buffer that actively increases the local chaperoning capacity of the nucleus. In contrast, cytoplasmic proteins, including translation factors and ribonucleoprotein granule components, exhibit increased aggregation under stress. Fluorescence microscopy confirmed that increased biochemical aggregation in density gradients reflects changes in assembly state *in vivo*. For many stress-induced aggregators, these changes are visible as spatial reorganisation in stressed cells, such as the formation of foci. Our findings suggest that cells use subcellular RNA distribution to tune the condensation and/or aggregation threshold of proteins in a compartment-specific manner.

## Introduction

Changing environmental conditions pose a continuous challenge to cells. To survive acute stresses such as heat, varying osmolarity of the surrounding medium, or nutrient limitation, cells must rapidly remodel their metabolism to maintain homeostasis under the new conditions. Gene expression is central to this adaptation, and regulation at every stage of RNA metabolism enables cells to mount an effective stress response (Bond, 2006). Many of the changes act to limit expression of the bulk transcriptome, while a small set of stress-induced transcripts gains privileged access to the gene expression machinery (Goswami et al., 2024; Seidler & Sträßer, 2024).

These changes in RNA metabolism are driven in large part by stress signalling. Kinase cascades convert stress signals into the coordinated phosphorylation of a large number of target proteins, including regulators of transcription, RNA processing, export and translation, and have long been recognised as a central layer of gene expression control (D. A. Smith et al., 2010; Toone & Jones, 1998). More recently, proteins prone to biomolecular condensate formation have emerged as a complementary layer of control (Dea & Pincus, 2024; Wallace et al., 2015). Many RNA-binding proteins (RBPs) are condensate-forming, positioning condensates at the centre of stress-induced changes to RNA metabolism (Alberti & Hyman, 2021; Roden & Gladfelter, 2021). Because demixing depends directly on physicochemical parameters such as temperature, pH, ionic strength, and macromolecular crowding, condensate-forming proteins can register a stress directly, without the need of additional upstream signalling input. The switch-like nature of this transition enables the appearance of a condensate to be read as a regulatory signal in itself (Riback et al., 2017; Villegas et al., 2022). The physiological impact of condensate-mediated regulation has been demonstrated in various systems: in mammalian cells, for example, the stress-responsive RBP ZFP36L2 facilitates the formation of condensates that degrade stress-associated mRNAs. This is a prerequisite for dedifferentiation, a reprogramming event that underlies tissue regeneration (Jiang et al., 2026).

Importantly, the two regulatory layers are not independent. Phosphorylation alters the charge and valency of the disordered regions that drive demixing, thereby promoting or suppressing the partitioning of an RBP into a condensate (Nosella & Forman-Kay, 2021). In turn, condensates recruit kinases, phosphatases, and chaperones, feeding back into the signalling networks that regulate condensate formation (Dea & Pincus, 2024; Söding et al., 2020; Wippich et al., 2013). Kinase signalling and condensation therefore act as an interconnected regulatory circuit.

This regulation affects gene expression at all stages. Activation of stress-specific transcription factors induces transcription of genes that harbour stress response elements in their promoters; one example is the heat-activated transcription regulator heat shock factor 1 (Hsf1) (Boy-Marcotte et al., 1999). Budding yeast Hsf1 is held in an inactive state by the chaperone Hsp70. The formation of heat-induced condensates sequesters Hsp70 away from Hsf1, leading to its activation and illustrating how condensate formation can be read out as a regulatory signal (Ali et al., 2023; Zheng et al., 2016). The co-occurring hyperphosphorylation of Hsf1 enhances transcriptional activity across systems (Lu et al., 2022; Zheng et al., 2016). The resulting stressspecific transcripts are efficiently exported from the nucleus, likely because Hsf1 interacts directly with the nuclear mRNA export receptor Mex67 (Zander et al., 2016). In contrast, export of most other mRNAs is inhibited during stress (Saavedra et al., 1996; Seidler & Sträßer, 2024).

Kinase signalling is important for this block, which suggests that phosphorylation of the export machinery plays a role (Carmody et al., 2010). Mechanistically, two changes have been observed. First, nuclear mRNPs components dissociate from the mRNA (Zander et al., 2016). Second, under some stress conditions, the DEAD-box ATPase Dbp5, which remodels mRNPs as they emerge from the nuclear pore, is depleted from the nuclear rim (Heinrich et al., 2024; Takemura et al., 2004). As a result, bulk mRNAs are largely retained in the nucleus.

Preferential expression of stress-specific transcripts is further enforced in the cytoplasm. Bulk translation is strongly reduced during stress (Goswami et al., 2024; Melamed et al., 2008; Simpson & Ashe, 2012; Williams & Rousseau, 2024), largely at the rate-limiting initiation step. Many stresses trigger phosphorylation of the alpha subunit of the translation initiation factor eIF2, which shuts down cap-dependent translation (Goswami et al., 2024; Simpson & Ashe, 2012; Williams & Rousseau, 2024). This translational repression coincides with the formation of RNA-protein condensates in the cytoplasm, most prominently stress granules and P-bodies, which are thought to act as storage sites for translationally repressed RNAs in yeast (Ripin & Parker, 2023). Beyond mRNAs, stress granules contain translation initiation factors, and a diverse set of RBPs and non-RBPs (Kedersha & Anderson, 2002; Kroschwald et al., 2015; Protter & Parker, 2016), whereas P-bodies are enriched in factors involved in mRNA turnover and translation repression (Kroschwald et al., 2015; Teixeira & Parker, 2007). Both types of granules are associated with endoplasmic reticulum (ER) tubules across organisms and are responsive to the secretory and translational capacity of the ER (Kilchert et al., 2010; Lee et al., 2020). How stress-specific transcripts escape translation repression is not understood in all its aspects, but condensation can again contribute (Hirose et al., 2023; Liu & Qian, 2014). In heatstressed budding yeast, sequestration in condensates of the RNA helicase Ded1, for example, which is required for efficient translation initiation of transcripts with highly structured 5’UTRs, selectively disadvantages these mRNAs and favours stress-induced transcripts with unstructured 5’-UTRs (Iserman et al., 2020). In other systems, both condensate evasion and preferential partitioning of newly synthesized transcripts to stress granules have been linked to their prioritised translation (Glauninger et al., 2025; J. Smith & Bartel, 2026). Condensate formation and translational reprogramming are therefore closely coupled. Condensates form as a consequence of reduced biosynthetic activity, and also influence which mRNAs are translated.

Taken together, stress reorganises RNA metabolism at every step, with many RBPs changing their RNA-dependent interactions and/or macromolecular assembly state. Proteome-wide RNA interactome methods have made one facet of this reorganisation, the remodelling of direct RNA-protein contacts, experimentally accessible. RNA interactome capture and poly(A)-independent approaches such as total RNA-associated proteome purification (TRAPP) and orthogonal organic phase separation (OOPS) use UV crosslinking followed by purification of crosslinked complexes to identify proteins in direct contact with RNA (Baltz et al., 2012; Castello et al., 2012; Kilchert, 2023; Queiroz et al., 2019; Shchepachev et al., 2019). Applied to stressed cells, these methods have revealed interactome changes that reflect the regulatory changes during cell adaptation outlined above. In budding yeast, for example, glucose starvation and heat stress cause translation initiation factors and ribosomal proteins clustered around the mRNA channel to lose contact with RNA (Bresson et al., 2020). A similar loss of initiation factor association with RNA occurs under oxidative stress (Matia-González et al., 2021), likely representing the interactome signature of translational shutdown. Conversely, TRAPP revealed strongly altered RNA association of the previously uncharacterised RBPs Pin4, Mrn1, Pbp2, and Nab6 following glucose starvation and heat stress (Bresson et al., 2020); Mrn1 and Nab6 were later shown to regulate levels of mRNAs encoding cell-wall-associated proteins (Bresson et al., 2023). Metabolic enzymes likewise alter their RNA association in a condition-dependent manner (Matia-González et al., 2021), suggesting that RNA binding by these proteins is itself regulated during stress adaptation.

These crosslinking approaches, however, detect direct RNA-protein contacts but not the macromolecular assembly states of the proteins involved. They therefore cannot capture the full spectrum of indirect, RNA-dependent associations found in larger RNP assemblies, where many components are recruited through proteinprotein interactions. Biochemical purification of stress granule cores from budding yeast and mammalian cells has demonstrated that condensate-derived RNP particles persist in cell lysates and be separated on the basis of their sedimentation behaviour (Demeshkina & Ferré-D’Amaré, 2025). To capture stress-induced changes in assembly state across the proteome, we profile the RNA-dependent sedimentation behaviour of proteins in cell lysates from stressed and unstressed conditions using the R-DeeP method. This density gradient-based approach employs quantitative mass spectrometry to monitor protein sedimentation in the presence and absence of RNA (Caudron-Herger et al., 2019, 2020; Rajagopal et al., 2025). We combine this with an analytical framework that utilizes the RNA-dependence index (RDI), which we previously developed to robustly compare sedimentation behaviour of proteins across gradients (Wäber et al., 2026). Specifically, we apply R-DeeP to *S. cerevisiae* exposed to heat or hyperosmotic stress (high salt). Both stresses alter the sedimentation profiles of numerous proteins, including shifts toward heavier fractions consistent with the incorporation into larger macromolecular assemblies, and increased aggregation in the absence of RNA.

This widespread exacerbation of aggregation upon RNA depletion provides proteome-wide support for the concept that RNA acts as a general chaperone, physically buffering proteins to prevent aberrant aggregation (Horowitz & Bardwell, 2016; Maharana et al., 2018). Crucially, our data suggest that cells may leverage this property during stress adaptation: the stress-induced retention of transcripts in the nucleus may actively increase nuclear chaperoning capacity and selectively shield the nuclear proteome from aggregation. Consistent with this hypothesis, many nuclear proteins escape stress-induced aggregation, but this selective protection is fully abolished by RNase treatment.

Across the proteome, shifts to higher molecular weight fractions correlated with the propensity of proteins to form microscopically visible condensates *in vivo*. These stress-responsive factors encompass translation regulators and established components of canonical RNP granules as well as proteins not previously described to associate with visible condensates or granules, pointing to a broader landscape of regulatory events driven by conditional protein condensation and spatial sequestration. Our data thus provide a proteome-wide resource for dissecting how macromolecular reorganization and spatial partitioning dictate cellular stress adaptation.

## Results

### R-DeeP profiling of heat- and salt-stressed *S. cerevisiae*

To examine how adaptation to severe stress remodels the RNA-dependent assembly state of the proteome, we selected two stress conditions for which stress-activated kinase signalling and the adaptive changes to the transcriptome have been studied extensively: high temperature and hyperosmotic stress (Galello et al., 2024). Cells were grown in rich medium at the optimal growth temperature of 30°C and then either shifted to 42°C (heat stress) or transferred to medium containing 1.25 M NaCl (salt stress) for one hour (Fig 1A, S1A), conditions that are associated with a strong nuclear mRNA export block (Seidler et al., 2026). Lysates from stressed or unstressed cells were analysed by R-DeeP (Caudron-Herger et al., 2019, 2020) as described previously (Wäber et al., 2026). In brief, lysates were subjected to density gradient ultracentrifugation with or without prior treatment with RNases, and the protein distribution across 25 fractions analysed by mass spectrometry (MS) (n = 3 biological replicates). Sedimentation profiles of all detected proteins under all conditions are accessible online: https://yeast-r-deep.computational.bio/ R-DeeP defines a protein as RNA-dependent if it sediments differently in the absence of RNA. This shift captures a spectrum of interactions, ranging from direct RNA binding and stoichiometric inclusion in defined ribonucleoprotein (RNP) complexes to looser associations with large, RNA-scaffolded assemblies. To quantify the difference between control and RNasetreated gradients, we calculated the RNA dependence index (RDI), a metric we introduced previously to allow robust cross-gradient comparisons (Wäber et al., 2026) (Figure 1A). As observed previously, proteins annotated as RNP components or RBPs had higher RDIs than other proteins in the untreated control, which was also true after stress treatment (Figure 1B and S1B) (Wäber et al., 2026). However, stress-specific patterns emerged: Heat stress broadened RDIs among RNP components. In contrast, salt stress shifted a subset of non-RBPs towards higher RDIs. This subset forms a visible shoulder in the RDI density plot, extending beyond our threshold established for RNA dependence (0.163) (Figure 1B, right panel) (Wäber et al., 2026).

**Figure 1:**
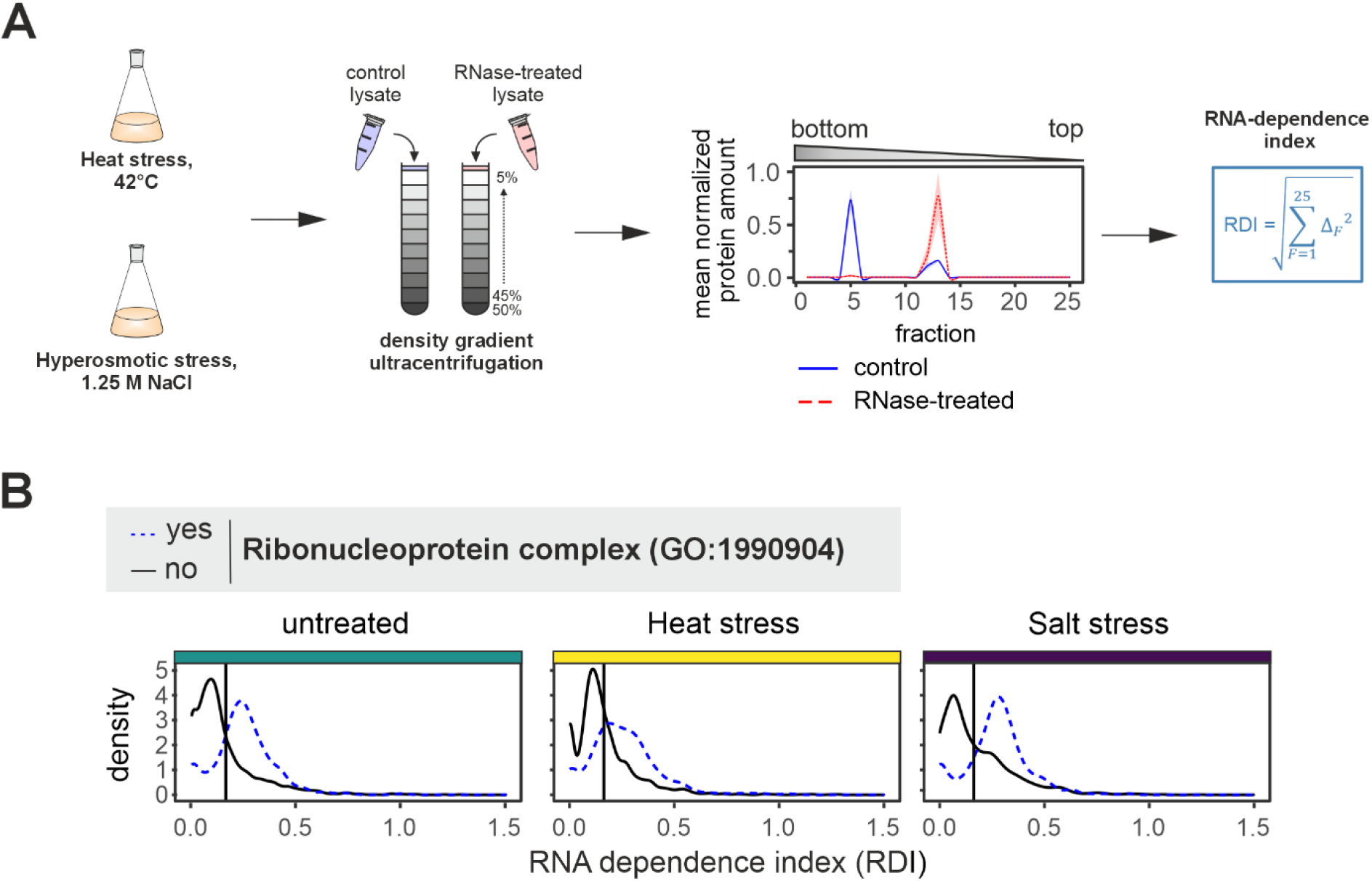
R-Deep profiling of heat- and salt-stressed *S. cerevisiae*. **A** Schematic of the R-DeeP workflow. For each protein in each condition tested, an RNA dependence index (RDI) was calculated as the Euclidean distance between sedimentation profiles in control and RNase-treated gradients. **B** Distribution of RDIs according to protein annotation “Ribonucleoprotein complex” (GO:1990904) for untreated, heat- and salt-stressed yeast cells. A threshold was applied to allow classification of proteins as RNA-dependent (RDI < 0.163) or -independent (RDI > 0.163), which is indicated by the vertical line. This threshold was previously determined based on the sedimentation behaviour of ribonucleoprotein complex components from *S. cerevisiae* in the absence of stress (Wäber et al., 2026).

### R-DeeP profiling reveals stress-specific changes to RNA-dependent protein sedimentation

To gain a more detailed perspective on how the two stresses alter the RNA-dependent assembly state of the proteome, we first performed principal component analysis (PCA) across all conditions. This revealed that proteins in untreated and salt-stressed samples behaved more similarly in both control and RNase-treated gradients, whereas heat stress produced a more extensive response (Figure 2A). To systematically identify groups of proteins whose RNA-dependent assembly state changes during stress adaptation, we hierarchically clustered all detected proteins based on their RDIs in untreated, heatstressed, or salt-stressed conditions (Figure S2A). These clusters broadly stratify the proteome by its baseline RNA dependence, allowing us to group proteins with distinct stress-responsive behaviour across both RBPs and unannotated proteins (Figure 2B and S2B and C). We identified a subset of proteins exhibiting a heatspecific increase in RNA dependence (cluster 4, with the mean RDI increasing to 0.32 from 0.12 in untreated cells) and another with increased RDI in response to both stresses (cluster 7, increasing from 0.3 to 0.46 under heat stress and 0.44 under salt stress, and enriched for proteins involved in translation; Figure 2C). Conversely, RNA dependence was specifically reduced after heat stress in several clusters, including clusters 8 and 9, which are both enriched for proteins involved in ribosome biogenesis and RNA processing (Figure 2C, D and S2D). Proteins associated with the different clusters are listed in Supplementary Table S1. Overall, changes to RNA-dependent sedimentation were more pronounced under heat stress, whereas sedimentation profiles under salt stress more closely resembled the untreated control.

**Figure 2:**
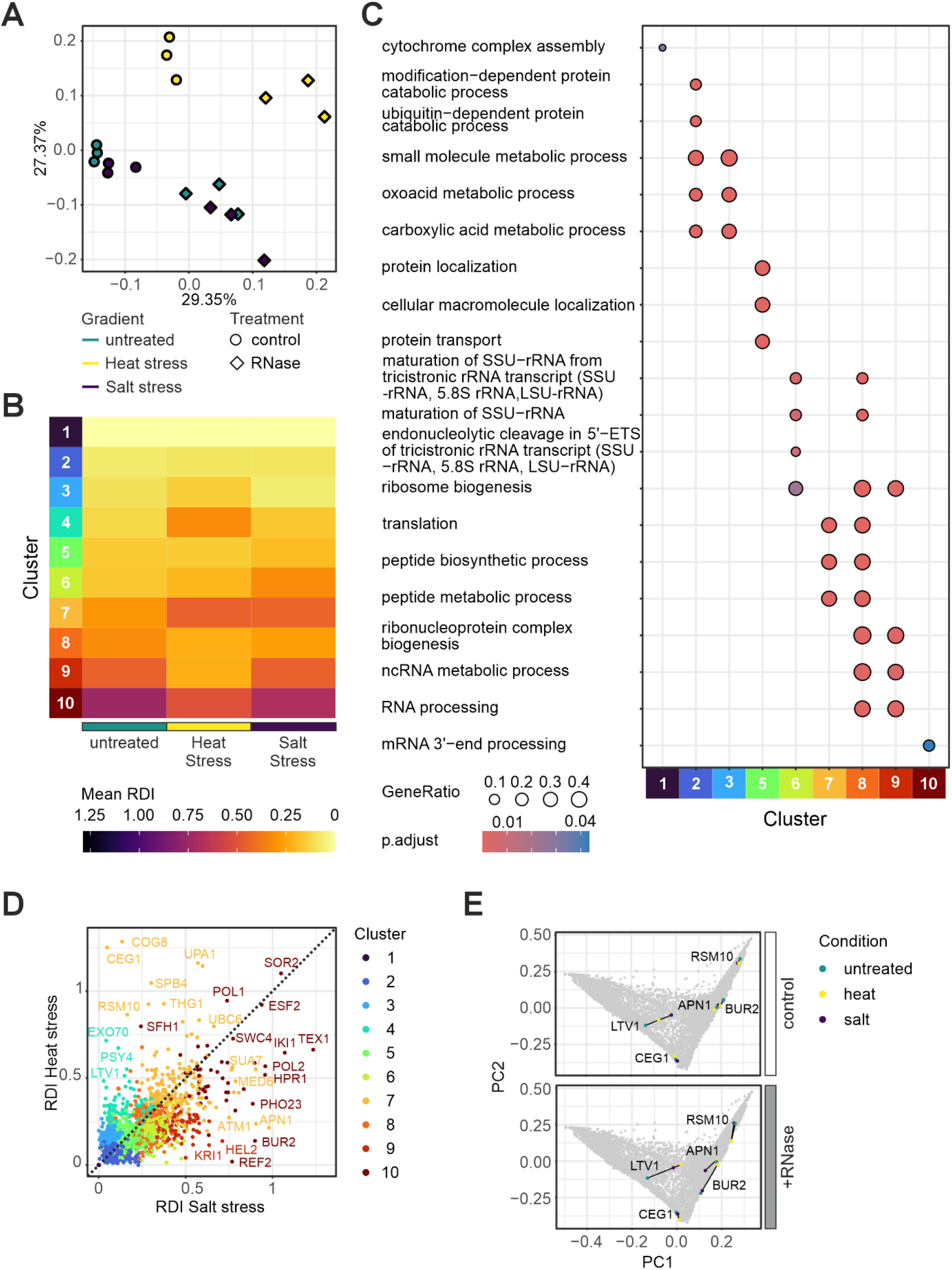
R-DeeP profiling reveals stress-specific changes to RNA-dependent protein sedimentation. **A** Principal component analysis (PCA) for the three biological replicates for all three conditions (untreated, heat and salt stress) with and without RNase treatment. The untreated, heat and salt stress gradients are coloured teal, yellow and purple, respectively. The control gradients are represented by circles and the RNase-treated gradients by a diamond. **B** Mean RNA dependence index (RDI) per cluster for proteins profiled across conditions. Proteins were grouped via hierarchical clustering (Fig S2A) and clusters sorted by their mean RDI in the untreated gradients. **C** Gene ontology (GO) term enrichment analysis (clusterProfiler) displaying the top three terms per cluster relative to the random distribution of terms across the full dataset. Node colour indicates significance (p-values adjusted for multiple testing using the Benjamini-Hochberg procedure), and circle size represents the gene ratio. **D** Pairwise comparisons of protein RDIs between heat- and salt-stressed conditions, with individual data points coloured according to their assigned clusters from panel B. **E** PCA for protein sedimentation profiles comparing all three conditions with and without treatment. To visualize the impact of stress, data points corresponding to representative proteins (selected from D) in control (top) or RNase-treated (bottom) gradients were highlighted. Point-to-point distance between conditions was calculated as indicated by the black lines.

Because the RDI serves as a summary metric for the magnitude of change rather than the direction of the shift, we subsequently examined the sedimentation profiles of proteins within specific clusters to identify typical patterns. Broadly speaking, we identified three notable patterns: In some cases, stress altered the amplitude of the shift without changing the peak location, resulting in a more intense signal loss or gain upon RNase treatment (clusters 4 and 7, Figures S3A and S3B). More strikingly, stress often induced a distinct shift towards higher molecular weight (MW) fractions (i.e., towards the bottom of the gradient), often resulting in aggregated signal in fraction #1, after RNase treatment (e.g., clusters 4, 6 to 9, Figure S3C and S3D). Conversely, a stress-specific reduction in RDI often occurred when a protein that in unstressed cells typically shifted towards lower MW fractions upon RNase treatment instead dispersed across the entire gradient or distributed across multiple smaller peaks (Figure S3C and S3D). However, stress treatment also caused more subtle changes in sedimentation behaviour, driving shifts of only a few fractions in opposite directions after RNase treatment, as observed in clusters 2 and 3 for proteins peaking around fraction 10 (Figure S3E). We conclude that stress adaptation triggers a complex physical reorganization of the RNA interactome, particularly under heat stress. Notably, stress-induced changes in RNA dependence are not uniform but reflect distinct patterns of altered sedimentation behaviour, such as redistribution into higher MW assemblies or dispersal of existing complexes.

To better account for these complexities, we applied principal component analysis (PCA) to the sedimentation profiles of each protein (Figure 2E). We then calculated the Euclidean distance between the resulting data points across the untreated, heat-stressed, and salt-stressed conditions, separately for control and RNase-treated gradients. Proteins whose distance in one stress condition to the other two conditions exceeded 0.1 were classified as stress-specific responders (Supplementary Tables S2 and S3). These responders showed two basic types of behaviour: Sedimentation of a subset of proteins was affected by stress treatment alone (e.g., Ltv1, Dsl1, and Tma19) (Figure 2E and S4A and B). More frequently, stress altered the responsiveness to RNase treatment, either sensitizing a protein to RNA removal and leading to a more pronounced shift after RNase treatment (e.g., Apn1), or rendering it more resistant (e.g., Bur2) (Figure 2E and S3D). The exonuclease Xrn1, for example, displayed identical sedimentation profiles in control gradients across all three conditions, and shifted to a lower-MW fraction following RNase treatment in untreated and salt-stressed yeast, yet became resistant to RNA removal after heat stress (Figure S4C and D). Looking at the heatspecific responders as a group, we found that they were enriched for GO terms associated with RNA-protein condensate formation relative to the full dataset (Figure S4E). Notably, as we had also observed for clusters 4 and 6 to 9 (Figure S3C and D), a large proportion of them shifted towards higher-MW fractions following heat exposure, which was exacerbated by RNase treatment (Figure S4F).

#### Stress and RNA depletion increase the propensity of proteins to form higher-order assemblies

The frequent shift of heat-responsive proteins towards higher-MW fractions raised the question of whether stress generally drives proteins into larger assemblies, and whether RNA buffers this effect. We therefore considered the direction of the shifts. When an RNA-protein complex dissociates following RNase treatment, its constituents typically shift towards the lighter top fractions of the sucrose gradient (positive shift). Shifts towards the heavier bottom fractions, in contrast, often reflect protein precipitation or aggregation (negative shift) (Caudron-Herger et al., 2019; Wäber et al., 2026). We therefore calculated the fraction offset between the signal maxima of two gradients, which gives the predominant direction of the shift for each protein (Figure 3A; note that for proteins that broadly disperse across the gradient the calculated offsets can be less conclusive).

**Figure 3:**
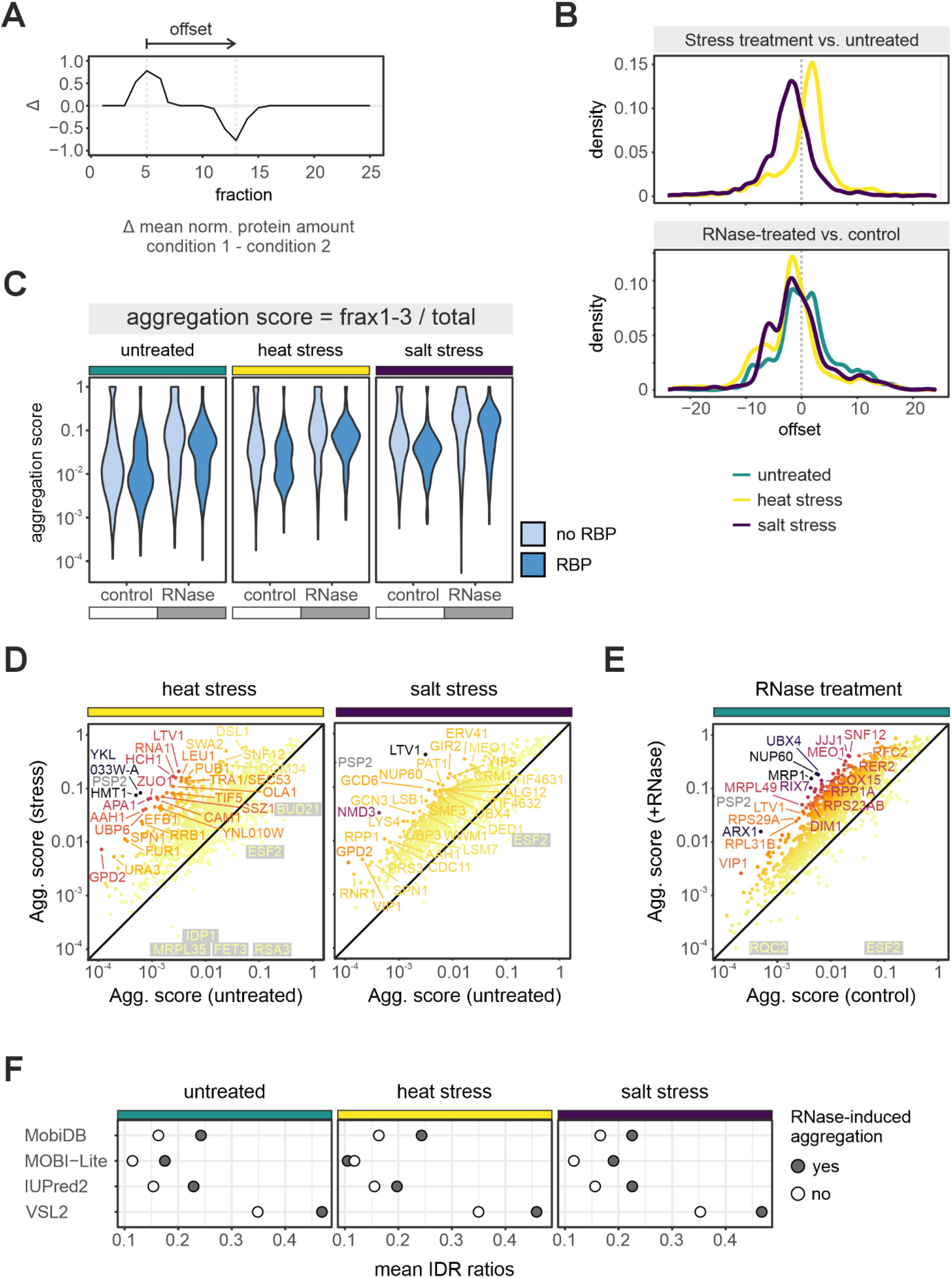
RNase and stress treatment induces protein aggregation in R-DeeP gradients. **A** Graphical representation of the offset calculation. **B** Density distribution of the calculated sedimentation offsets. The top panel shows the shifts induced by stress alone (relative to the untreated control), while the bottom panel depicts the RNA-dependent shifts induced by RNase treatment across all three conditions. Negative offsets indicate a shift towards heavier, higher molecular weight fractions. **C** Aggregation score distribution with and without RNase treatment across the three conditions grouped by whether proteins are annotated as RNA binding. Aggregation scores reflect the proportion of protein signal in the three bottom-most fractions of the gradient. Non-aggregating proteins were not included. **D** Comparison of protein aggregation scores following heat stress (left) and salt stress (right) relative to the untreated control. Colouring represents fold-change of aggregation score after stress relative to untreated. **E** Comparison of protein aggregation scores following RNase treatment relative to the control (untreated cells). Colouring represents fold-change of aggregation score after RNase treatment relative to control. **F** Comparison of median IDR content between proteins that do or do not undergo RNase treatment-induced aggregation, defined as 10-fold increase in aggregation score resulting in at least 5 % of total protein being present in the three bottom-most fractions; values represent consensus scores generated using MobiDB, MOBI-Lite, IUPred2, and VSL2, designed to predict and score intrinsic disorder in protein structures, and retrieved from(FrenchPacheco et al., 2022).

First, we assessed whether stress treatment alone drives proteins towards heavier fractions relative to the untreated control (Figure 3B, top panel). Salt stress indeed caused a strong tendency towards negative offsets (area under the curve [AUC] below 0 = 0.73). Although many strongly heat-responsive proteins shifted towards the bottom fractions (e.g., clusters 4, 6 and 9, Figure S3C, S4F), these proteins are a minority. Across the proteome, heat stress mostly led to positive offsets, with only about a third of proteins displaying negative offsets. This likely reflects the lower stability of many protein complexes at elevated temperature; as destabilized complexes dissociate into their individual components, they sediment less deeply into the gradient and shift towards lighter fractions.

Next, we evaluated the combined effect of stress and RNA depletion. RNase treatment induced both positive and negative offsets across all conditions (Figure 3B, bottom panel). However, following stress exposure, RNA depletion drove the majority of shifting proteins to negative offsets (AUC below 0: untreated = 0.47; heat stress = 0.68; salt stress = 0.60). This marked redistribution into heavier fractions suggests that RNA broadly protects proteins from precipitating or forming higher-order aggregates in stressed cells, and that this protection is lost when RNA is removed.

Because our offset calculation considers the shift of a protein’s main peak, it can miss aggregation that affects only a fraction of the molecules of a given protein. To capture this, we calculated an aggregation score for each protein, defined as the proportion of its total signal detected in the three bottom-most fractions of the gradient (fractions 1-3). Most proteins had a low basal aggregation score (below 0.1) in unstressed control gradients (Figure S5A), but RBPs were more prone to aggregate than non-RBPs (64% and 50% non-zero aggregation score, respectively). Proteins that constitutively aggregate strongly (scoring above 0.5 across all conditions) were largely membrane-associated, consistent with either co-sedimentation with membrane remnants or aggregation of exposed hydrophobic surfaces (Figure S5B). We also identified five RBPs with high constitutive aggregation scores, Whi2, Whi3, Edc3, Rom1, and Sgd1; of these, both the prion-like protein Whi3 and the P-body component Edc3 have been reported to phase separate (Currie et al., 2023; Peskett et al., 2025), whereas the other three have not been linked to condensation or formation of pathological aggregates.

Outside this group, aggregation scores increased markedly both following stress and in response to RNase treatment (Figure 3C, D, E and S5C). Notably, heat stress raised aggregation scores to a similar extent as salt stress, even though it shifted the main peak of most proteins towards positive offsets. This is consistent with aggregation affecting only a subpopulation of each protein, which the offset does not capture. Importantly, RNase treatment increased aggregation scores across all conditions, including in unstressed cells, but affected stressed cells most strongly (Figure 3D, E and S5C). Thus, the aggregation score further supports the offset analysis, suggesting that RNA broadly protects proteins against aggregation, and that this protection becomes more important under stress. Because previous work has implicated RNA as a critical chaperone that prevents aggregation of IDR-containing RBPs in particular (Horowitz & Bardwell, 2016; Maharana et al., 2018; Park et al., 2024), we assessed the structural disorder of proteins with increased aggregation after RNase treatment (> 10-fold increase with at least 5% of total protein being present in the three bottommost fractions). Indeed, based on multiple prediction algorithms (French-Pacheco et al., 2022), aggregating proteins possessed higher mean structural disorder than the rest of the dataset (Figure 3F). This robust correlation provides proteome-wide support for the concept that RNA acts as a general chaperone, physically buffering IDR-containing proteins to prevent their aberrant precipitation.

### Stress-induced aggregation is compartmentalized and buffered by nuclear RNA

If RNA functions as a general chaperone for IDR-containing proteins, its subcellular distribution should determine its local protective buffering capacity. During environmental stresses, including heat and salt stress, eukaryotic cells retain mRNAs in the nucleus, leading to their accumulation (Saavedra et al., 1996; Seidler & Sträßer, 2024). This export block is classically viewed as a regulatory step to suppress bulk translation. We hypothesized that it may simultaneously serve a biophysical function. By actively increasing the RNA chaperoning capacity of the nucleus, it could protect the nuclear proteome from stress-induced aggregation. Consistent with previous reports (Saavedra et al., 1996; Seidler et al., 2026), poly(A)+ RNA accumulated in the nucleus under both stresses (Figure 4A and B). Ultrastructural analysis showed that RNA was not evenly distributed across nuclear subcompartments, with poly(A)+ RNA enriched in regions that stained poorly with DAPI under both stresses, most likely the nucleolus (Figure 4C). Under salt stress, the RNA signal was in addition punctate and almost exclusively found in discrete foci, whose composition is currently unknown (Figure 4C).

**Figure 4:**
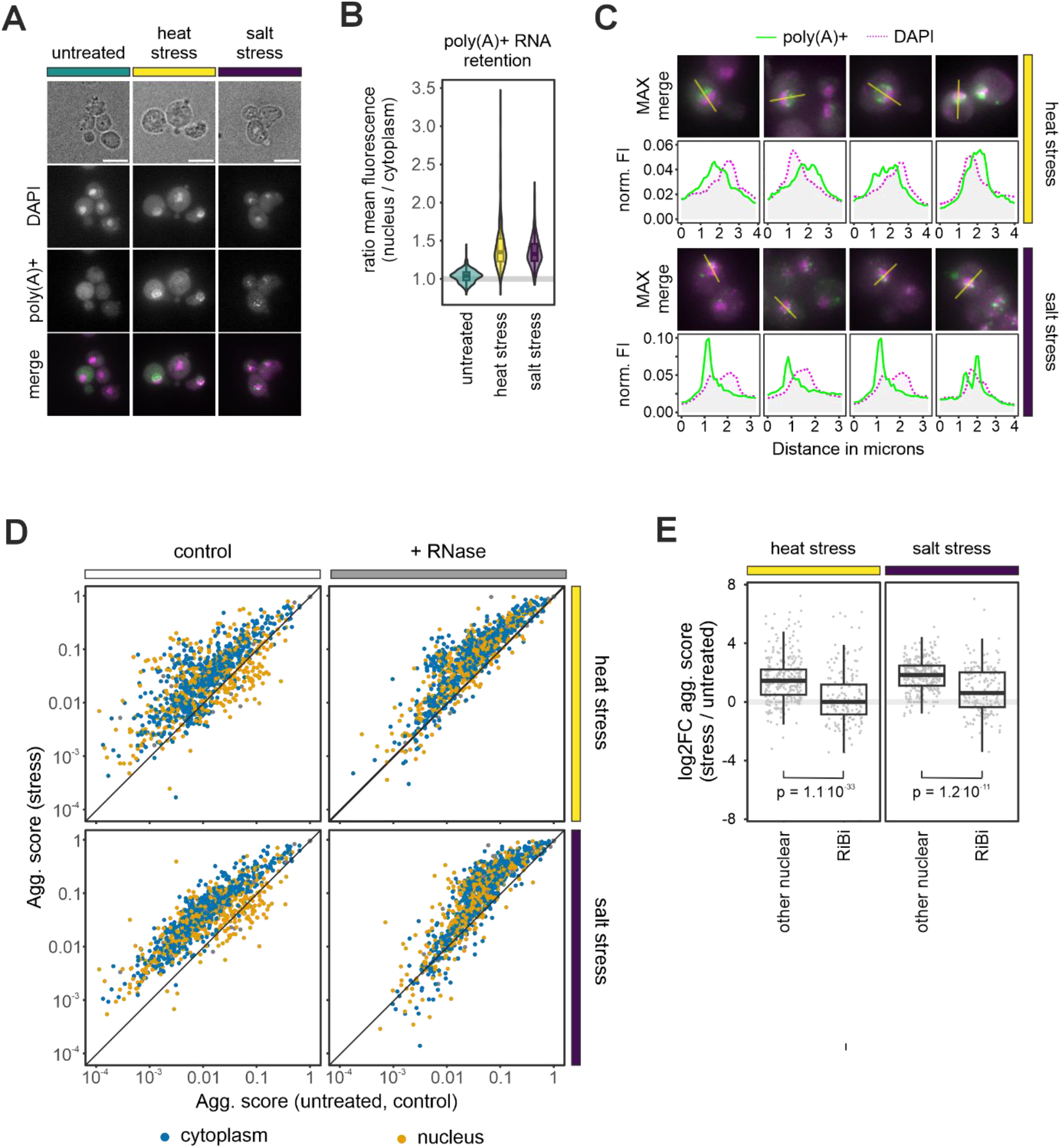
Stress-induced aggregation is compartmentalized. **A** Fluorescence in-situ hybridisation (FISH) against poly(A)+ RNA using a Cy3-labelled oligo-d(T) probe and DAPI to stain the DNA showing a single Z plane. The merged channel shows poly(A)+ RNA in green and DAPI in magenta. Scale bar corresponds to 5 microns. **B** Quantitation of oligo-d(T) FISH signal as shown in A. Measurements were performed on average intensity Z-projections for the nucleus and cytoplasm and the ratio of mean nuclear over cytoplasmic fluorescence intensity calculated for each cell. (n = 4 and 3 independent experiments for heat stress and salt stress, respectively, with > 64 cells counted for each). **C** poly(A)+ FISH and DAPI signal (maximal projection, MAX) after heat stress (top) and salt stress (bottom). Fluorescence intensity (FI) profiles were measured along the yellow lines, with FI normalized to total area under the curve. **D** Comparison of protein aggregation scores following heat stress (top) and salt stress (bottom) in the presence (left) and absence (right) of RNA relative to the untreated control, with proteins annotated with GO cellular compartment “nucleus” (GO:0005634) in orange, all others in blue. Proteins for which no annotations could be retrieved are shown in grey. **E** Log2 fold change increase in aggregation score (log2FC agg. score) following stress for proteins involved in ribosome biogenesis (GO:0042254) and all other nuclear proteins in the dataset. The displayed p-values for the pairwise comparisons were calculated using a twosided Wilcoxon rank sum test.

To test whether nuclear-retained RNA protects nuclear proteins, we compared the aggregation scores of nuclear and cytoplasmic proteins between conditions. Consistent with the buffering hypothesis, compartment-specific analysis showed that cytoplasmic proteins displayed a higher propensity to aggregate under both heat and salt stress, whereas nuclear proteins were partially protected (Figure 4D, left panels). Crucially, when we degraded RNA in the stressed lysates, this compartment-specific protection was abolished (Figure 4D, right panels), demonstrating that it depends on RNA. Furthermore, analysis of intrinsic disorder content showed that nuclear proteins shielded from stress-induced aggregation possessed significantly higher median IDR scores compared to other nuclear proteins (Figure S5D), which matches the properties expected of RNA chaperone clients. Functionally, they were enriched for proteins annotated in ribosome biogenesis (RiBi), a process that takes place in the nucleolus. Consistent with this, RiBi proteins on average showed a significantly smaller increase in aggregation score under stress than other nuclear proteins (Figure 4E). The proteins protected from aggregation are therefore found in the compartment where RNA accumulates under stress. Together, these data strongly suggest that the stress-induced nuclear retention of RNA may act as a compartmentspecific buffering system that dynamically redistributes chaperoning capacity to the nucleus during stress.

### Stress drives aggregation of IDR-rich proteins, including translation regulators

A natural consequence of a redistribution of chaperoning capacity to the nucleus would be a relative loss of protection from aggregation in the cytoplasm, which we indeed observed in our stress-treated samples (Figure 4D). We next asked whether this compartmentalized decrease in chaperoning activity might serve a regulatory purpose by promoting the selective aggregation of specific functional networks. To systematically classify proteins with strong aggregation in response to stress alone, we applied the same stringent threshold as before: Any protein with at least a 10-fold increase in aggregation score resulting in at least 5% of total protein being present in the three bottom-most fractions was classified as aggregating in a stress-dependent manner. This identified 71 and 35 heatand salt-specific aggregators, respectively, with a core set of 10 proteins aggregating under both conditions (Figure 5A; Supplementary Table S4). Stressspecific aggregators were associated with diverse starting distributions in the untreated gradients, and RNase treatment generally exacerbated their aggregation further (Figure 5B). Notably, these stress-specific aggregators included nuclear as well as cytoplasmic proteins and showed only a minor overlap with proteins previously reported to aggregate during arsenite stress or stationary phase, whereas 18 of our heat stress aggregators were shared with a dataset of proteins that aggregated in response to high heat (46°C) (Figure S5E) (Ibstedt et al., 2014; Narayanaswamy et al., 2009; Wallace et al., 2015). As IDR-rich proteins were enriched both among proteins that aggregate after RNase treatment (Figure 3F) and among nuclear proteins shielded from stress-induced aggregation (Figure S5D), we asked whether proteins that aggregate under stress share this feature. Indeed, stress-specific aggregators had a higher predicted mean structural disorder compared to the rest of the dataset (Figure 5C). GO analysis further showed that stress-induced aggregators are associated with cytoplasmic assemblies linked to RNA metabolism, such as cytoplasmic RNPs and stress granules, as well as with membranous structures (Figure 5D). This fits the view that RNP assemblies are frequently driven by IDRs that provide multivalent interaction sites (Protter & Parker, 2016). The enriched processes and functions mirrored the GO component results: stress-induced aggregators were enriched for roles in translation and its regulation, particularly translation initiation (Figure 5D). This is consistent, because translation initiation factors (TIFs) are core components of stress granules (Ivanov et al., 2019). Importantly, the trend was not limited to proteins that met our stringent aggregator criteria. In response to salt as well as heat stress, proteins known to partition into cytoplasmic stress granules or P-bodies, as well as proteins annotated as translation regulators, had higher median aggregation scores compared to the untreated control and other proteins, respectively (Figure 5E and S6A, B). TIFs in particular showed increased aggregation under both stresses, but individual complexes responded with different sensitivity; while translation initiation factor 2 (eIF2) subunits were affected similarly by both stresses, eIF3 and eIF5 responded more strongly to heat, whereas eIF4F components responded more strongly to salt (Figure S6C). Aggregation of the translation initiation machinery into stress granules during stresses that cause a global shutdown of bulk translation, including heat stress, is well described and has been proposed to contribute to the control of translation (Cherkasov et al., 2013; Grousl et al., 2009; Ivanov et al., 2019). Our data show that salt stress drives the same behaviour. These observations are compatible with a model of reduced cytoplasmic buffering. RNA depletion is unlikely to be the only driver, however. Under stress, initiation factors aggregated significantly more strongly than after RNase treatment alone (Figure S6D), clearly showing that other mechanisms are involved; reduced cytoplasmic buffering may, however, promote or facilitate TIF aggregation.

**Figure 5:**
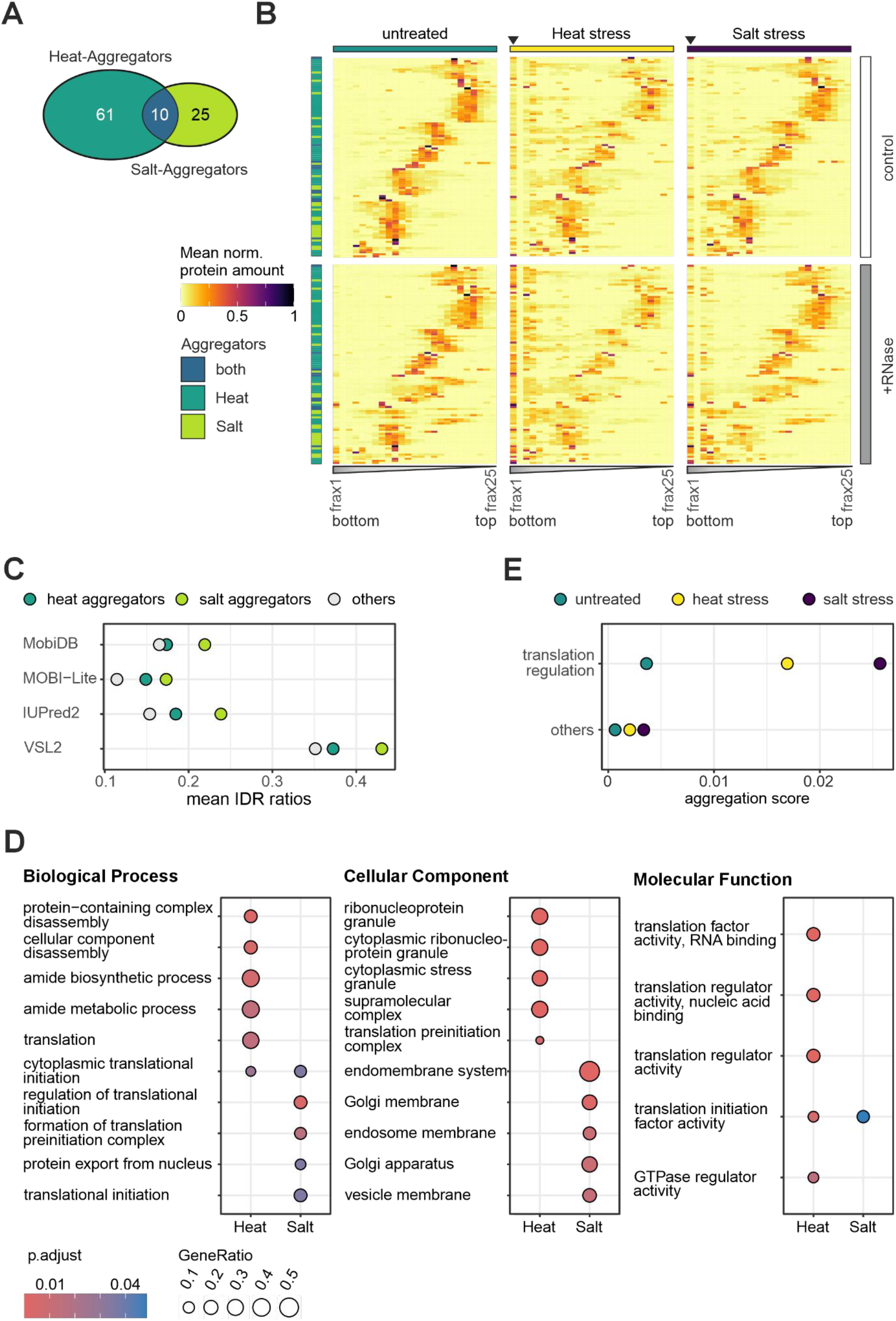
Stress drives aggregation of IDR-rich proteins, including translation regulators. **A** Identification of stress-specific aggregators for heat and salt stress, defined as proteins exhibiting at least a 10-fold increase in aggregation score relative to the untreated control, comprising at least 5% of their total protein amount. **B** Sedimentation profiles of stress-specific aggregators in control (upper panel) and RNase-treated (lower panel) gradients across all three conditions. Colouring to the right of the graph indicates aggregation only under heat stress (dark green), only under salt stress (light green), or under both conditions (blue). The black triangle marks the bottom-most fraction of the stress gradients. **C** Mean IDR ratios for heatand salt-specific aggregators compared to the remaining proteins detected in the dataset; values represent consensus scores generated using MobiDB, MOBI-Lite, IUPred2, and VSL2, designed to predict and score intrinsic disorder in protein structures, which were retrieved from (French-Pacheco et al., 2022). **D** GO term enrichment analysis of heat stress- and salt stress-specific aggregators relative to the random distribution of terms across the full dataset. Node colour and size indicate significance and gene ratio, respectively. The top five cellular component GO terms per cluster are shown. **E** Comparison of aggregation scores of proteins annotated with “translation regulation” in untreated, heat-stressed, and salt-stressed proteins compared to the remaining proteins in the dataset.

### Many stress-specific aggregators form microscopically visual cellular foci

To confirm that stress-induced biochemical aggregation in gradients reflects the formation of cellular condensates or aggregates *in vivo*, we monitored the subcellular distribution of selected stress-specific aggregators from the cytoplasm and the nucleus by fluorescence microscopy using genomically GFP-tagged strains. We focused our microscopic validation on salt stress, because several heat-induced aggregators in our dataset, including Arc1, Gus1, Mes1, Ola1, Pub1 and components of eIF3, have already been shown to form condensates or aggregates in cells during heat stress (Cherkasov et al., 2013; Desroches Altamirano et al., 2024; Wallace et al., 2015). We selected salt stress-responsive aggregators with diverse functions, including translation regulation (Ded1, Gcd6, Gir2, Tif4631), RNA decay (Pat1, Xrn1), signalling (Ett1), nuclear export (Crm1), and ribosome biogenesis (Ltv1, selected as an exception to the general protection of this class) (Figure 6A). Some of these proteins were previously described to partition to canonical cytoplasmic granules under various types of stresses (P-bodies: Pat1, Xrn1; stress granules: Tif4631; shared: Ded1). The majority of stress-induced aggregators formed microscopically detectable foci in response to salt treatment, although their specific appearance varied (Figure 6B). For canonical P-body and stress granule components, the GFP signal was largely diffuse in the cytoplasm of unstressed cells, with Ded1 appearing slightly patchier. Following one hour of salt stress, all of these markers coalesced into a distinctly granular pattern. While shorter (15 min) salt stress has been reported to induce P-bodies but not stress granules (Buchan et al., 2008; Kilchert et al., 2010), our prolonged 1h stress captured condensation of Tif4631 (eIF4G), which has been characterised as a stress granule component (Figure 6B) (Hoyle et al., 2007; Mutlu et al., 2019). This is consistent with the established sequential order of P-body and stress granule formation: Across systems, P-bodies have been reported to form first and nucleate stress granule formation (Buchan et al., 2008; Lee et al., 2020; Majerciak et al., 2023). We also observed stress-induced granular GFP signal for the translation regulator Gir2, which was not previously described as a granule component, although its condensation phenotype was relatively weak. In contrast, the eIF2B subunit Gcd6 exhibited a distinctive change in localization. In unstressed cells, Gcd6 frequently localized to a large, constitutively present cytoplasmic body, as has been previously described (Campbell et al., 2005); upon salt stress, this large body fully dispersed into a distributed granular pattern that more closely resembled the canonical P-body or stress granule markers (Figure 6B).

**Figure 6:**
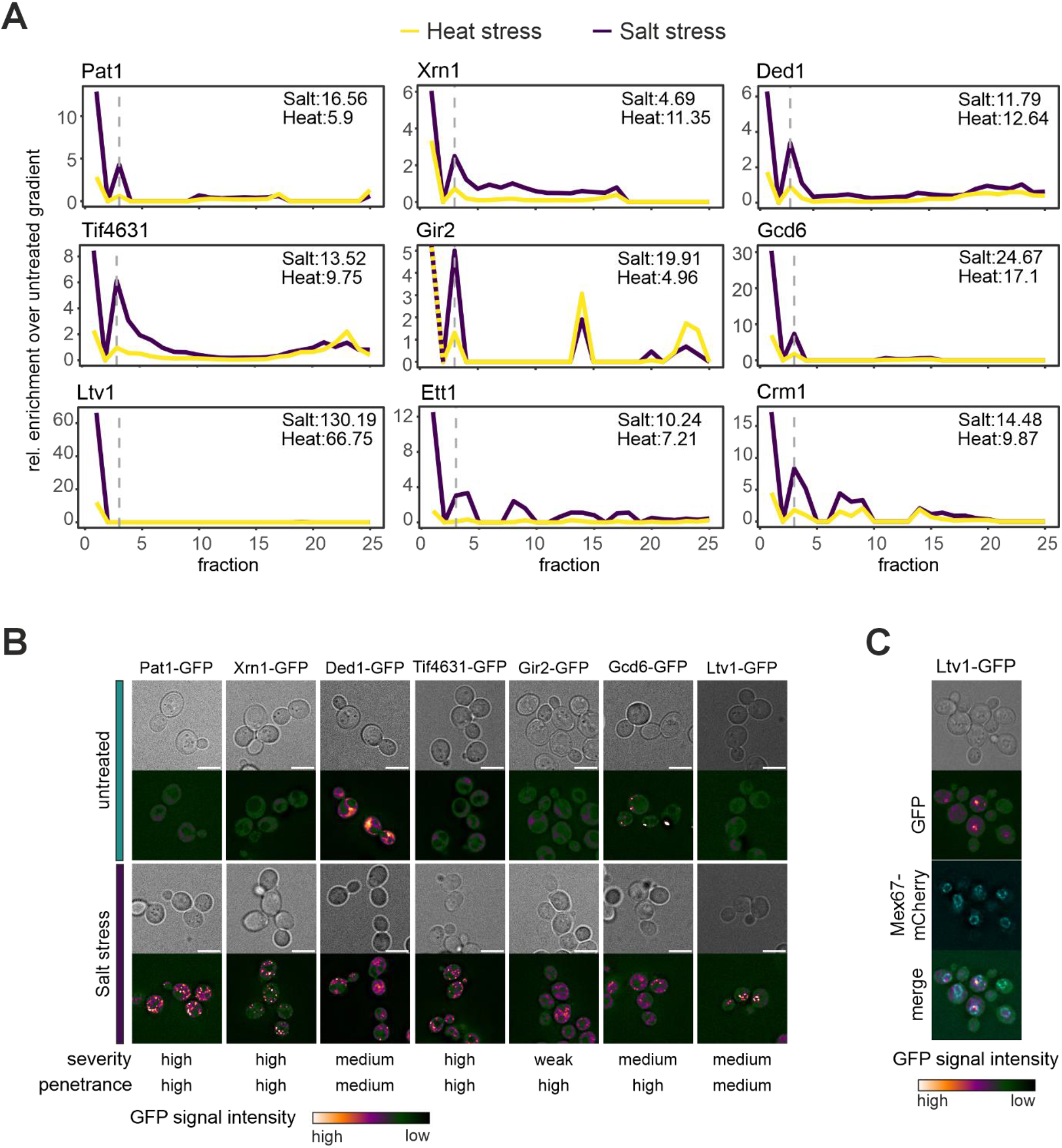
Proteins with high aggregation scores form microscopically visual foci under stress. **A** Relative enrichment of selected proteins in heat (yellow) and salt stress (purple) gradients relative to the untreated control. The calculated fold-change in aggregation score is indicated in the top right of each plot. A dashed gray line demarcates fractions 1 to 3, which correspond to high-molecular-weight aggregates (Note: The enrichment value for Gir2 in the bottom fractions is theoretically infinite, as no Gir2 was detected in fractions 1-2 of the untreated gradient). **B** Representative live-cell fluorescence microscopy images of the GFP-tagged proteins analysed in A, comparing untreated cells with cells subjected to salt stress (n = 3 biological replicates). The relative severity and penetrance of the observed condensation phenotypes are summarized below the images. Across all microscopy panels, GFP signal intensity is visualized using a false-colour scale (Blue Orange icb), as indicated at the bottom. Contrast and brightness settings were kept constant between untreated and salt-stressed conditions for each protein. Scale bars correspond to 5 microns. **C** Salt stress-induced foci formation of Ltv1-GFP as in B, co-expressed with the nuclear envelope marker Mex67-mCherry (shown in cyan) to confirm spatial localization (n = 3 biological replicates).

Foci formation was not restricted to the cytoplasm. Ltv1 is a RiBi factor, a class that is on average protected from stress-induced aggregation yet showed a severe aggregation phenotype in response to salt stress (130-fold increase in aggregation score; Figure 6A). Ltv1 was diffuse in the absence of stress but formed prominent condensates upon salt exposure that were limited to a smaller volume of the cell (Figure 6B). By co-expressing the nuclear envelope marker Mex67-mCherry, we confirmed that these stress-induced Ltv1 foci are constrained within the nucleus (Figure 6C). Proteins with highly specific localization in unstressed cells were more difficult to evaluate. The nuclear protein Ett1, which was classified as a “superaggregator” during high heat (Wallace et al., 2015), and the export factor Crm1 localized to the nucleoplasm and the nuclear rim, respectively, under both unstressed and salt-stressed conditions (Figure S6E). Their signal became noticeably brighter and potentially more condensed after salt stress. However, we could not distinguish whether this is a condensation event that is qualitatively different from simple accumulation at its constitutive localization site, for example due to increased molecular crowding. These two cases therefore remain inconclusive. Of the nine salt-induced aggregators we tested, seven showed stress-induced foci or a clear change in granular pattern, including Gir2 and Ltv1, which were not previously known to partition to condensates. Together with the previously documented *in vivo* aggregation of heat-induced aggregators from our dataset (Cherkasov et al., 2013; Wallace et al., 2015), these results show that Increased aggregation in R-DeeP gradients reports on changes in protein assembly state inside cells, under both heat and salt stress. Because gradients also capture other transitions in protein assembly state, individual candidates still require microscopic confirmation. Regardless, these data show that the dataset can be used to discover stress-induced condensation events and to prioritize proteins for cell biological follow-up experiments.

## Discussion

In this study, we apply R-DeeP to map RNA-dependent changes in protein assembly state in *S. cerevisiae* during stress. Following both heat and salt stress, we observed distinct changes in the RNA dependence of factors involved in RNA metabolism, particularly for proteins that regulate translation and RNP granule dynamics. The most prominent response was a shift of proteins into high-MW fractions, indicative of increased aggregation. For most proteins tested by microscopy, this biochemical precipitation correlated with the formation of microscopically visible cellular foci (Figure 6). Aggregation in density gradients therefore appears to be a reliable indicator of stress-induced reorganisation of proteins into larger assemblies *in vivo*.

The stress-induced aggregation we observe likely captures a spectrum of high-MW assemblies, ranging from functional biomolecular condensates to irreversible protein aggregates. Cellular aggregates typically consist of misfolded proteins associating via exposed hydrophobic residues, or highly stable amyloid-like folds, which are often sequestered into dedicated compartments for chaperone-mediated clearance (Alberti & Hyman, 2021; Lippi & Krisko, 2024). Conversely, functional condensates, such as stress granules and P-bodies, are dynamic structures formed by multivalent protein-protein, RNA-protein and RNA-RNA interactions (Ripin & Parker, 2023). Because IDRs frequently drive the assembly of both functional condensates and pathological aggregates, the two structures share overlapping biophysical properties (Ripin & Parker, 2022; Visser et al., 2024). Indeed, we found that both stress granule-associated proteins and prion-like proteins such as Whi3 aggregate in the control gradients (Figure S6). However, the sedimentation profiles of aggregating proteins vary widely from protein to protein, even for proteins associated with the same condensates, and are distributed heterogeneously over fractions #1 to #4. This suggests that our method captures intermediate assemblies or subcomplexes of varying sizes, consistent with findings that different proteins exchange at different rates and are present in intact cellular granules with varying stoichiometries (Xing et al., 2020). It is also consistent with liquid-to-solid transitions of RNP condensates that have been observed in response to stress (Guillén-Boixet et al., 2020; Kroschwald et al., 2015; Yang et al., 2020). Aggregation scores are therefore a simplified readout of assembly state, which does not only reflect the size of cellular granules. This is illustrated by the behaviour of Gcd6, which forms a large cytoplasmic body in unstressed cells that is not associated with aggregation in the gradient (Campbell et al., 2005) (Figure 6A and B). Upon salt stress, this body disperses into a more granular pattern that is associated with significant aggregation (Figure 6B). The nature of these stress-induced granules is unclear.

RNase treatment broadly exacerbated protein aggregation in our gradients, including that of canonical RNP granule components, and the proteins most affected were enriched for IDRs (Figure 3F). Our observations strongly support the model of RNA chaperoning or buffering, which posits that high intracellular RNA concentrations keep IDR-containing RBPs soluble by occupying their binding sites and preventing pathological aggregation (Horowitz & Bardwell, 2016; Maharana et al., 2018). A related mechanism has recently been proposed for long 3’-untranslated regions (3’-UTRs), which may act as chaperones that enable co-translational folding of proteins with high IDR content (Luo et al., 2026). Our lysate-based approach has a precedent in *E. coli*, where Park and Seong used quantitative mass spectrometry to analyse the contribution of RNA to proteome solubility and were able to relate it, on the protein side, to IDR content and isoelectric point (Park et al., 2024). The immediate, exacerbated aggregation we observe in RNase-treated gradients is therefore likely a consequence of IDRs being left unchaperoned after RNA removal. However, we cannot exclude that RNase digestion also disrupts RNPs in ways that expose aggregation-prone surfaces independently of IDR content. Whether features of the RNA also influence chaperoning efficiency is less clear, but there is circumstantial evidence. A recent integrative profiling study identified a class of condensation-prone RNAs (“smOOPs”) that show low sequence complexity and are heavily bound by RBPs (Klobučar et al., 2026). The study addressed condensation and not RNA buffering; however, it is conceivable that RNAs that have a strong capacity to engage with IDR-rich RBPs are particularly effective chaperones. If so, RNA buffering would depend not only on how much RNA is present in a compartment but also on which RNAs are present. Whether the RNAs that are retained in the nucleus during stress are enriched in smOOPlike or other features that favour chaperoning remains an open question.

Irrespective of which RNAs are involved, our data suggest a general principle: that cells use RNA as a broadly acting chaperone under stress, and that global RNA retention or redistribution of RNA can act as a protective mechanism. Aggregation-proneness is not a general feature of the proteome but a property of a specific subset of proteins for which RNA promotes solubility. Most proteins show no aggregation in untreated cells or after RNase treatment (Figure S5A). Proteins that aggregate strongly after RNase treatment almost always already show measurable aggregation in untreated controls; conversely, with very few exceptions, proteins with measurable aggregation in untreated control gradients aggregate more strongly after RNase treatment (Figure 3E). The set of aggregationprone proteins and the set of RNA-buffered proteins are therefore nearly identical.

In stressed yeast cells, nuclear RNA retention leads to a massive accumulation of transcripts in the nucleus, most prominently in DAPI-poor regions that likely correspond to the nucleolus (Figure 4C). Nuclear proteins are partially protected from aggregation, and this protection is lost after RNase treatment. This shielding effect is most pronounced for RiBi factors, which function in the nucleolus (Figure 4E). This suggests that nucleolar proteins are the most likely target of prioritised protection in yeast. We therefore propose that RNA retention increases the local chaperoning capacity of the nucleus and shields nuclear proteins from stress-induced aggregation. Cytoplasmic proteins, which are comparatively less protected, aggregate more readily. This shift in protective capacity could explain why translation regulators, which are largely cytoplasmic, are preferentially affected, and would link the subcellular redistribution of RNA to the shutdown of bulk protein synthesis. Translation initiation is a point where many forms of control converge. It is the rate-limiting step of translation and targeted by several stress-responsive mechanisms, including phosphorylation of eIF2a and sensitivity to ATP depletion (Bexley et al., 2025; Wek et al., 2023). Aggregation of TIFs, promoted by compartment-specific RNA depletion, may add further input at this regulatory hub; however, the relative contribution of this mechanism under different conditions remains to be tested, and the model is so far correlative. Beyond the oligo(dT)-FISH experiment (Figure 4B), we have no quantitative data measuring RNA concentrations in each compartment. The increased nucleolar and reduced cytoplasmic protection are inferred from the relative aggregation of nucleolar and cytoplasmic proteins, respectively. Together with the preferential protection of RiBi factors, this suggests that the ribosome biogenesis machinery is either particularly vulnerable to damage or particularly worth protecting.

Nuclear RNA retention appears to be an active process in yeast. It depends on signalling through the cell wall integrity kinase Slt2, and nuclear poly(A)+ accumulation in response to heat stress is lost in its absence (Carmody et al., 2010). Intriguingly, *Δslt2* cells still produce heat shock proteins, indicating that retention is not required for their efficient expression. This suggests that retention serves another purpose, possibly the chaperoning function we propose here.

Stress-induced nuclear RNA accumulation is not limited to budding yeast. It is also observed in stressed plant cells and after virus infections in human cells (Seidler & Sträßer, 2024). In viral infection, activation of RNase L blocks mRNA export, which has been suggested to act as an antiviral mechanism that limits host and viral protein synthesis (Burke et al., 2021). Where RNA accumulates may nevertheless differ between systems and conditions. In *S. pombe*, nucleolar proteins accumulate in nucleolar rings upon heat shock, which also contain poly(A)+ RNA (Aydin et al., 2024; Gallardo et al., 2020). In mammalian cells, poly(A)+ RNA accumulates in nuclear speckles (McIntyre et al., 2025). We suggest that these are variations on one theme in which stress-induced RNA retention concentrates a non-specific chaperone where it is needed most.

Nuclear RNA retention does not, however, shield all nuclear proteins from aggregation. Ltv1, a ribosome biogenesis factor required for the maturation and export of the 40S subunit (Remines et al., 2026; Seiser et al., 2006), is an example of this. Although RiBi factors are, on average, shielded from aggregation, Ltv1 forms prominent nuclear foci and aggregates strongly under salt stress (Figure 6A-C). It was previously identified as a heat-induced “superaggregator” (Wallace et al., 2015), but in our data its behaviour depends on the stressor. Under heat stress, where toxicity is thought to be predominantly driven by protein unfolding, RNase treatment exacerbated Ltv1 aggregation (Figure 2D), consistent with RNA keeping it soluble. Under salt stress, aggregation was independent of RNA, arguing against buffering under these conditions. Increased molecular crowding may instead drive the aggregation (Figure 6A, B). Given the role of Ltv1 in 40S maturation, its condensation could pause ribosome biogenesis during stress. Whether this is an adaptive response is unclear, because deleting *LTV1* reduces ribosome levels and sensitises cells to osmotic and oxidative stress (Loar et al., 2004; Remines et al., 2026). Whether Ltv1 foci contain RNA also remains to be tested. However, there is precedent for retained RNA being associated with nuclear assemblies. The mRNA export factor Nab2 forms nuclear foci during heat and glucose starvation (Carmody et al., 2010; Heinrich et al., 2024). Its aggregation depends on RNA but not on its IDR and correlates with mRNA retention in a dose-dependent manner during glucose starvation (Heinrich et al., 2024). Retained RNA may therefore have more than one role. It can shield some nuclear proteins from aggregation, as we propose for RiBi factors, but it can scaffold or nucleate nuclear assemblies of others.

The idea of RNA as a non-specific, exploitable chaperone also connects to other fields. In neurodegenerative disease, aggregation of IDRrich, prion-like RBPs such as FUS and TDP-43 is central to pathology. RNA can keep such proteins soluble and prevent their aggregation (Maharana et al., 2018; Mann et al., 2019; Silva et al., 2022). Both proteins are mostly nuclear, where RNA is abundant, and their aggregation is typically associated with mislocalisation to the cytoplasm, where RNA is less abundant. This parallels the shift in chaperoning capacity that we propose under stress. If the subcellular distribution of RNA influences proteostasis, its redistribution, whether stress-induced or disease-related, could decide which proteins aggregate. Whether this principle extends beyond the systems and conditions studied here remains to be tested.

## Materials and Methods

### Yeast strains and strain generation

All yeast strains used in this study are listed in Supplementary Table 5. Plasmids and oligos used for strain generation are listed in Supplementary Tables 6 and 7, respectively.

### Stress treatment and lysate preparation

A 2L main culture was grown to an OD_600_ of 0.8 in YDP (1% yeast extract, 2% peptone, 2% glucose). For stress treatment, 500 ml cultures were mixed with 500 ml 54°C warm medium or with 500 ml YPD containing 2.5 M NaCl to reach the final temperature of 42°C or concentration of 1.25 M NaCl, respectively. The cultures were grown for 1h at 42°C for heat and 30°C for salt stress. Cells were harvested by centrifugation at 3600 rpm (Avanti JXN-26 with JLA-8.1) for 3 min at RT and washed with 5 ml lysis buffer (25 mM Tris, pH 7.5, 100 mM KCl, 0.02% (v/v) Triton-X; added freshly: 0.5 mM DTT, 1 μl/10 ml buffer protease inhibitor, and 1 × PhosStop (Roche)). Cells were lysed in 500 µl lysis buffer with 0.3 g glass beads for four cycles (20 s, 6m/s) using a Fastprep-24 5G with 60 sec on ice between cycles.

### R-DeeP density gradient analysis

R-Deep gradient preparation, centrifugation, fractionation, sample preparation for MS, MS measurements and protein identification were performed as described before (Wäber et al., 2026). In brief, a sucrose gradient was prepared ranging from 50 to 5% sucrose by subsequent layering and freezing of the gradient. The gradient was removed from the freezer and thawed 6 h prior to centrifugation. The protein amount in the lysate was determined using a BCA assay and RNase digest was performed for 1h at 4°C for the RNase-treated samples. Control samples were incubated with addition of an equivalent volume of buffer. After, the lysate was carefully layered on top of the gradient and centrifuged for 18 h 30000 rpm at 4°C. Bottom-to-top fractionation was performed using an Äkta Star. Proteins were precipitated using E-TCA precipitation. For MS-analysis, proteins were digested with trypsin and each sample TMT-labelled using TMTpro-18-plex (Thermo Fisher Scientific). Reagent set was used as follows: 129N-Heatctrl1, 130C-Heat-ctrl2, 131C-Heat-ctrl3, 130NHeat-rnase1, 131N-Heat-rnase2, 132N-Heatrnase3, 132C-Hyperosmotic-ctrl1, 133C-Hyperosmotic-ctrl2, 134C-Hyperosmotic-ctrl3, 133N-Hyperosmotic-rnase1, 134N-Hyperosmoticrnase2 and 135N-Hyperosmotic-rnase3. After labelling, all samples were combined by fraction number and proteins identified by liquid chromatography and tandem mass spectrometry (LC-MS/MS/MS).

### Data analysis

R scripts used for data analysis are available on github (https://github.com/Kilchertlab/R-DeeP). GO term annotations were retrieved from Ensembl using biomaRt (Durinck et al., 2009; Kinsella et al., 2011) or using org.Sc.sgd.db (Carlson, 2017) and AnnotationDbi (Pagès & Carlson, 2026). GO term analysis was performed using ClusterProfiler (Wu et al., 2021) and org.Sc.sgd.db. Hierarchical clustering was performed using pheatmap (Kolde, 2025) and visualized using ggplot2 using the viridis colour scales (Garnier et al., 2024; Wickham, 2016). Spine plots were generated using geom_xspline() function from the ggalt package (Rudis et al., 2016), correlation heatmaps were generated using ComplexHeatmap (Gu et al., 2016), PCA was performed using the pcromp function from the R stats package, and visualized using ggplot2 (Wickham, 2016). The area under the curve was calculated with the MESS package (Ekstrøm, 2025). Predictions of structural disorder for the yeast proteome were retrieved from (French-Pacheco et al., 2022).

### RNA FISH

Cells were grown to an OO_600_ of 0.8 and stress induced as described above. Cells were fixed with 3.7% formaldehyde for at RT for 40 min. Cells were washed and the cell wall permeabilised using 100 µg Zymolyase T100 (Carl Roth) for 30 min at 37°C. Cells were incubated in prehybridization buffer (50 % formamide, 10 % dextran sulphate, 125 μg/ml *E. coli* tRNA, 500 μg/ml herring sperm DNA, 4 x SSC, 0.02 % polyvinyl pyrrolidone, 0.02 % BSA, 0.02 % Ficoll-400) for 1h at 37°C before adding 0.75 µl of 1 pmol/μl oligo-(dT)50-Cy3 for incubation overnight. Cells were washed with 0.5x SSC and mounted on poly-lysine-coated cover slips in Roti Mount FluorCare DAPI (Carl Roth). Z-stack images were acquired on a Deltavision Ul-tra High-Resolution Microscope (Cytiva) using AcquireUltra (1.2.2) and deconvolved with soft-WoRx (7.2.1) using default settings. Image crops and line profiles were generated with Fiji (ImageJ 1.53t) (Schindelin et al., 2012). For quantitation, the mean poly(A)+ RNA signal per compartment was determined by segmenting nuclei and cells on the DAPI and the oligo-(dT) signal, respectively, with the cytoplasmic compartment defined as the cell body without the nucleus. Nuclear to cytoplasmic RNA ratios for each cell were calculated in R and plotted with ggplot2 (Wickham, 2016).

### Live-cell imaging

Cells were grown in rich medium and NaCl added to a final concentration of 1.25M. After 1h, cells from 400 µl were pelleted and resuspended in 25 µl clear medium (EMMG with or without 1.25 M NaCl). 3 µl were mounted on poly-lysine coated coverslips. Z-stack images were acquired on a Deltavision Ultra High-Resolution Microscope (Cytiva) using AcquireUltra (1.2.2) and deconvolved with softWoRx (7.2.1) using default settings. Image crops were generated with Fiji (ImageJ 1.53t) (Schindelin et al., 2012).

## Data accessibility

The MS proteomics data have been deposited to the ProteomeXchange Consortium via the RIDE partner repository (Deutsch et al., 2023) with the dataset identifier PXD058607. Gradient profiles can be accessed at https://yeast-r-deep.computational.bio/, where the complete processed datasets are also available for download. A generalized Snakemake workflow for normalization and analysis of R-DeeP fractionation data is available at Zenodo (https://doi.org/10.5281/zenodo.21297290).

## Acknowledgments

We thank Laura Henke-Schulz for the generation of the Mex67-mCherry expression vector. This work was supported by the Deutsche Forschungsgemeinschaft (DFG) via the Emmy Noether Programme (KI1657/2-1 to C.K.) and the graduate training group GRK2355 (325443116 to A.G., C.K. and K.S.); the European Research Council (ERC Consolidator Grant, mRNP-PackArt to K.S.); the Cardio-Pulmonary Institute (CPI), EXC 2026 (to K.S.); and the de.NBI Cloud within the German Network for Bioinformatics Infrastructure (de.NBI) and ELIXIR-DE (W-de.NBI-001, Wde.NBI-004, W-de.NBI-008, W-de.NBI-010, W-de.NBI-013, W-de.NBI-014, W-de.NBI-016, W-de.NBI-022). Gemini Pro 3.1 (Google) and Claude Sonnet 5.5 (Anthropic) were used to support writing.

