## Supplementary Figures S1-6 and Supplementary Tables 5-7 for "Proteome-wide profiling reveals compartment-specific RNA buffering of protein aggregation during stress"

### Supplementary Data

#### Supplementary Tables

##### Supplementary Table 5: Strains used in this study

| ### | name | genotype | source |
|---|---|---|---|
| Y1 | wild-type | *ade2–1, trp1–1, leu2–3, his3–11, ura3–52, lys+, can1–100* | |
| YPCK1094 | GCD6-GFP | *ade2–1, trp1–1, leu2–3, his3–11, ura3–52, lys+, can1–100, gcd6-GFP::kanMX* | This study |
| YPCK1095 | TIF4631-GFP | *ade2–1, trp1–1, leu2–3, his3–11, ura3–52, lys+, can1–100, tif4631-GFP::kanMX* | This study |
| YPCK1096 | CRM1-GFP | *ade2–1, trp1–1, leu2–3, his3–11, ura3–52, lys+, can1–100, crm1-GFP::kanMX* | This study |
| YPCK1097 | LTV1-GFP | *ade2–1, trp1–1, leu2–3, his3–11, ura3–52, lys+, can1–100, ltv1-GFP::kanMX* | This study |
| YPCK1099 | ETT1-GFP | *ade2–1, trp1–1, leu2–3, his3–11, ura3–52, lys+, can1–100, ett1-GFP::kanMX* | This study |
| YPCK1100 | DED1-GFP | *ade2–1, trp1–1, leu2–3, his3–11, ura3–52, lys+, can1–100, ded1-GFP::kanMX* | This study |
| YPCK1101 | Xrn1-GFP | *ade2–1, trp1–1, leu2–3, his3–11, ura3–52, lys+, can1–100, xrn1-GFP::TRP1* | This study |
| YPCK1115 | PAT1-GFP | *ade2–1, trp1–1, leu2–3, his3–11, ura3–52, lys+, can1–100, pat1-GFP::kanMX* | This study |
| YPCK1116 | GIR2-GFP | *ade2–1, trp1–1, leu2–3, his3–11, ura3–52, lys+, can1–100, gir2-GFP::kanMX* | This study |
| YPCK1130 | Ltv1-GFP Mex67-mCherry | *ade2–1, trp1–1, leu2–3, his3–11, ura3–52, lys+, can1–100, ltv1-GFP::kanMX* + pRS314-Mex67-mCherry | This study |

**Supplementary Table 6: Plasmids used in this study**

| ### | name | source |
|---|---|---|
| KS2218 | pRS314-Mex67-mCherry | Laura Henke-Schulze |
| | pYM26 (yeGFP-KITRP1) | Janke et al., Yeast 2004 |
| CK009 | pCRII-TOPO-L3/L6-rrp6-GFP-KanMX | Aydin et al., Nat Commun. 2024 |

**Supplementary Table 7: Oligos used in this study**

| Name | Sequence | Purpose |
|---|---|---|
| Xrn1_S2 | tatactattaaagtaacctcgaatatacttcgtttttagtcgtatgttctaATCGATGAATTCGAGCTCG | C-terminal tagging |
| Xrn1_S3 | tcacaaagcaatgctgctgaccgtgataataaaaaagacgaatctactCGTACGCTGCAGGTCGAC | C-terminal tagging |
| Xrn1_colony | AGCGAATGATGTTGCAGATGT | Strain validation |
| ScPAT1-L4-rc | GTTATGGGGTTGGTGTATCGCGATGGTGAAATATCAGAACTAAAGcggatccccgggttaattaa | C-terminal tagging |
| ScPAT1-L5-rc | aaaaaatacatgcgtaagtacattaaaattacaggaaaaatcTTAatcgatgaattcgagctcgaattacaggaaaaatcTTAatcgatgaattcgagctcg | C-terminal tagging |
| ScPAT1-tagtest-F | CCACTATTAACGAGGCGGAG | Strain validation |
| ScGIR2-L4-rc | GATGACGTAGCCAAGGGACTTGCCAAGACCGAAATAGCAAATCAAcggatccccgggttaattaa | C-terminal tagging |
| ScGIR2-L5-rc | ccatgcagaagaaaatacttaaatacttaagtacacaagtatTTAatcgatgaattcgagctcg | C-terminal tagging |
| ScGIR2-tagtest-F | AAGGCTCATCATGGGAAGCT | Strain validation |
| ScGCD6-L4-rc | GAGTGGTTACAGAATGCTGACGAAGAATCTTCCTCAGAAGAGGAAcggatccccgggttaattaa | C-terminal tagging |
| ScGCD6-L5-rc | ttttatggtaatacatgattatgcagggtactgcgtggacttTTAatcgatgaattcgagctcg | C-terminal tagging |
| ScGCD6-tagtest-F | AGTGCATTGGTTTCTCTATACGA | Strain validation |
| ScTIF4631-L4-rc | ACAAATATGTTCAGTGCATTAATGGGAGAAAGTGATGACGAAGAGcggatccccgggttaattaa | C-terminal tagging |

| | | |
|---|---|---|
| ScTIF4631-L5-rc | tatccaagtgacattttcgatacttaacatgatctattcatgTTAatcgatgaattcgagctcg | C-terminal tagging |
| ScTIF4631-tagtest-F | GCACGACAAGAACTGGAACA | Strain validation |
| ScCRM1-L4-rc | GCTGCCAAGATTGGTGGGTTATTAAAACCTTCCGAACTTGATGATcggatccccgggttaattaa | C-terminal tagging |
| ScCRM1-L5-rc | ggaaaaaatattggaaatttaaagaatgatacgccaccgcctCTAatcgatgaattcgagctcg | C-terminal tagging |
| ScCRM1-tagtest-F | GGTGCAAATCAAAGAAGTCGG | Strain validation |
| ScLTV1-L4-rc | AAGAAACTTGAGAAGGTCACCAATACACTAAGCAGCTTAAAATTTcggatccccgggttaattaa | C-terminal tagging |
| ScLTV1-L5-rc | agtacttgtaatgtaggtgctttctcatctcattctactcctCTAatcgatgaattcgagctcg | C-terminal tagging |
| ScLTV1-tagtest-F | GAGAGTGGTGGCCGTAAACT | Strain validation |
| ScETT1-L4-rc | TCTCATGGTAAATTTCAAGATGTCCTGGACAATTTCTTACAAGGCcggatccccgggttaattaa | C-terminal tagging |
| ScETT1-L5-rc | acagaaggagtgagtaaaaataaaaaagaagcgtggatgcgcTCAatcgatgaattcgagctcg | C-terminal tagging |
| ScETT1-tagtest-F | TAGCGACAAGGAAGCCAAGA | Strain validation |
| ScDED1-L4-rc | TCTGGCTGGGGTAACAGCGGTGGTTCAAACAACTCTTCTTGGTGGcggatccccgggttaattaa | C-terminal tagging |
| ScDED1-L5-rc | tagagcagaaaacgaagaatcctcaccctagtttgtctgaaaTCAatcgatgaattcgagctcg | C-terminal tagging |
| ScDED1-tagtest-F | ACGCTATGATGAGTGCTCCA | Strain validation |

#### Supplementary Figures

### Figure S1

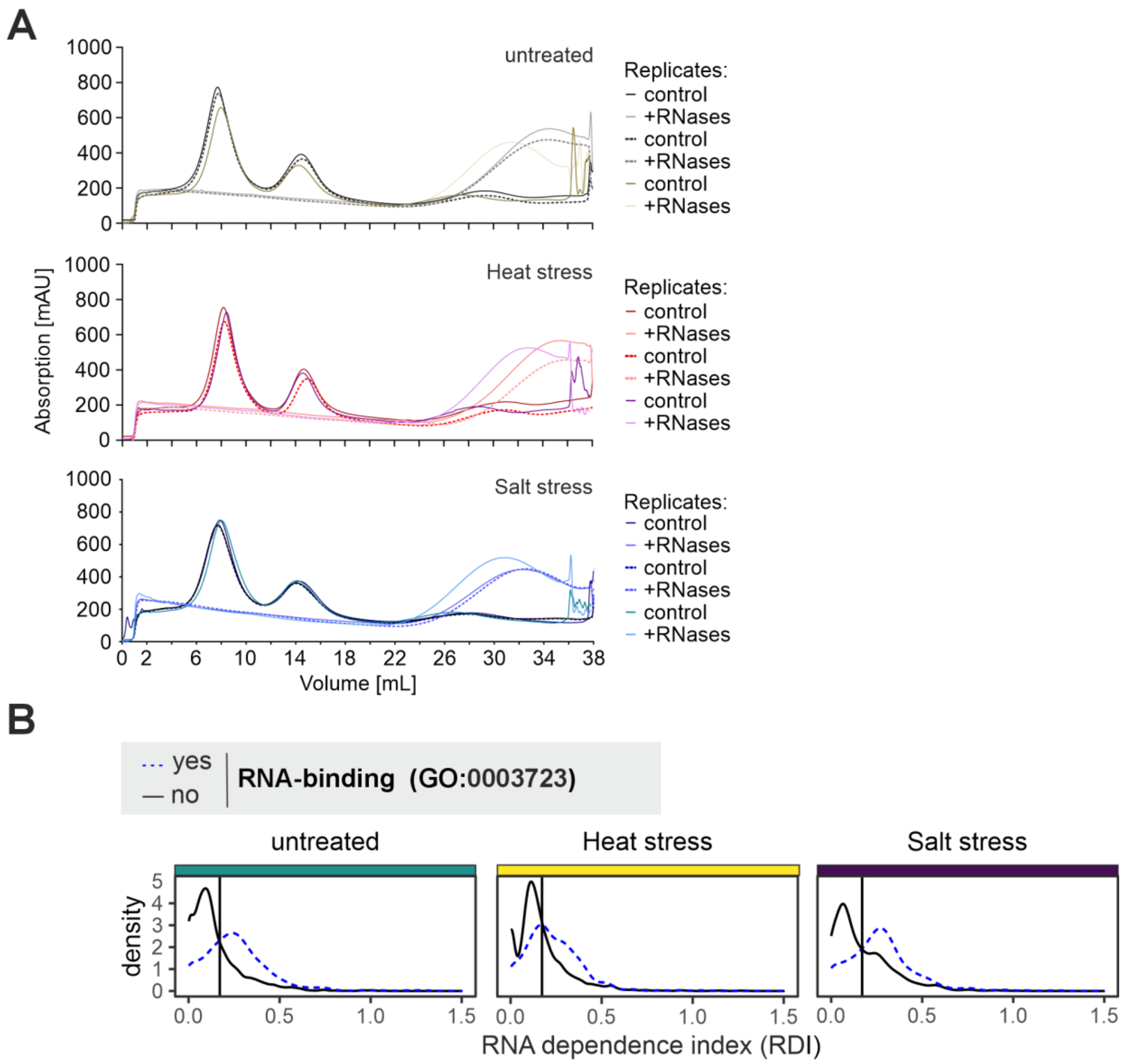


**Figure S1: A** UV absorption profile at 280 nm recorded during the sucrose gradient fractionation of control and RNase-treated lysates. Profiles are shown for all three biological replicates across the untreated, heat-stressed, and salt-stressed conditions. **B** Distribution of RNA dependence index (RDI) values for untreated, heat-stressed, and salt-stressed cells, grouped by annotation “RNA-binding” (GO:0003723). The vertical dashed line indicates the RDI threshold previously established to classify proteins as RNA-dependent (RDI > 0.163) or RNA-independent (RDI < 0.163), based on the baseline sedimentation behaviour of RNP components in unstressed *S. cerevisiae* (Wäber et al 2026).

### Figure S2

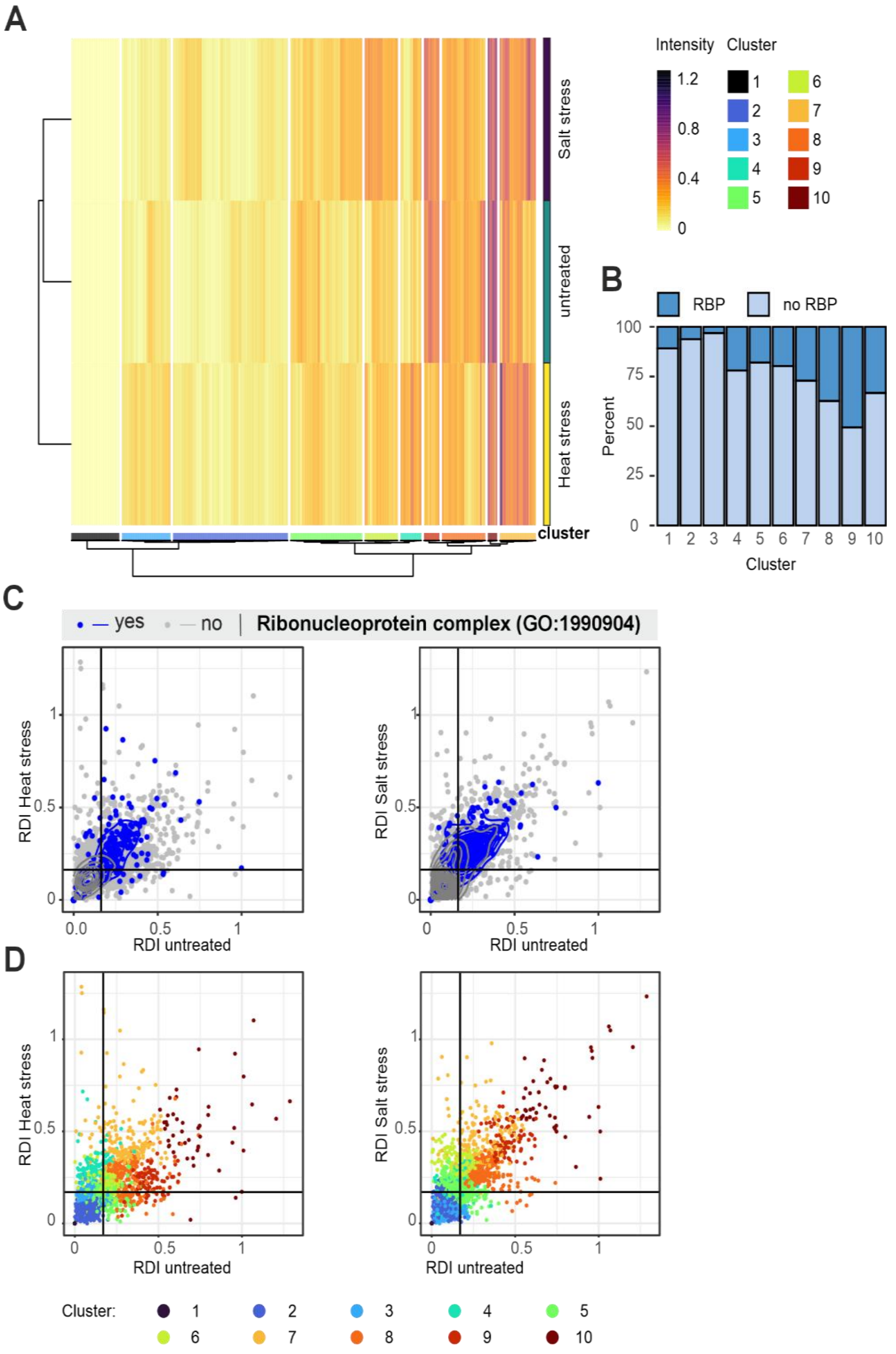


**Figure S2: A** Hierarchical clustering of proteins detected across all three conditions based on their RDIs. **B** Percentage of annotated RNA-binding proteins (RBPs) within each identified cluster. **C** Pairwise comparisons of protein RDIs between stressed (heat or salt) and untreated conditions. Proteins annotated as ‘ribonucleoprotein complex’ components (GO: 1990904) are highlighted in blue, while unannotated proteins are shown in grey. Contours indicate 2d density estimates for the populations. **D** Pairwise RDI comparison as shown in C, with individual data points coloured according to their assigned clusters from panel A.

### Figure S3

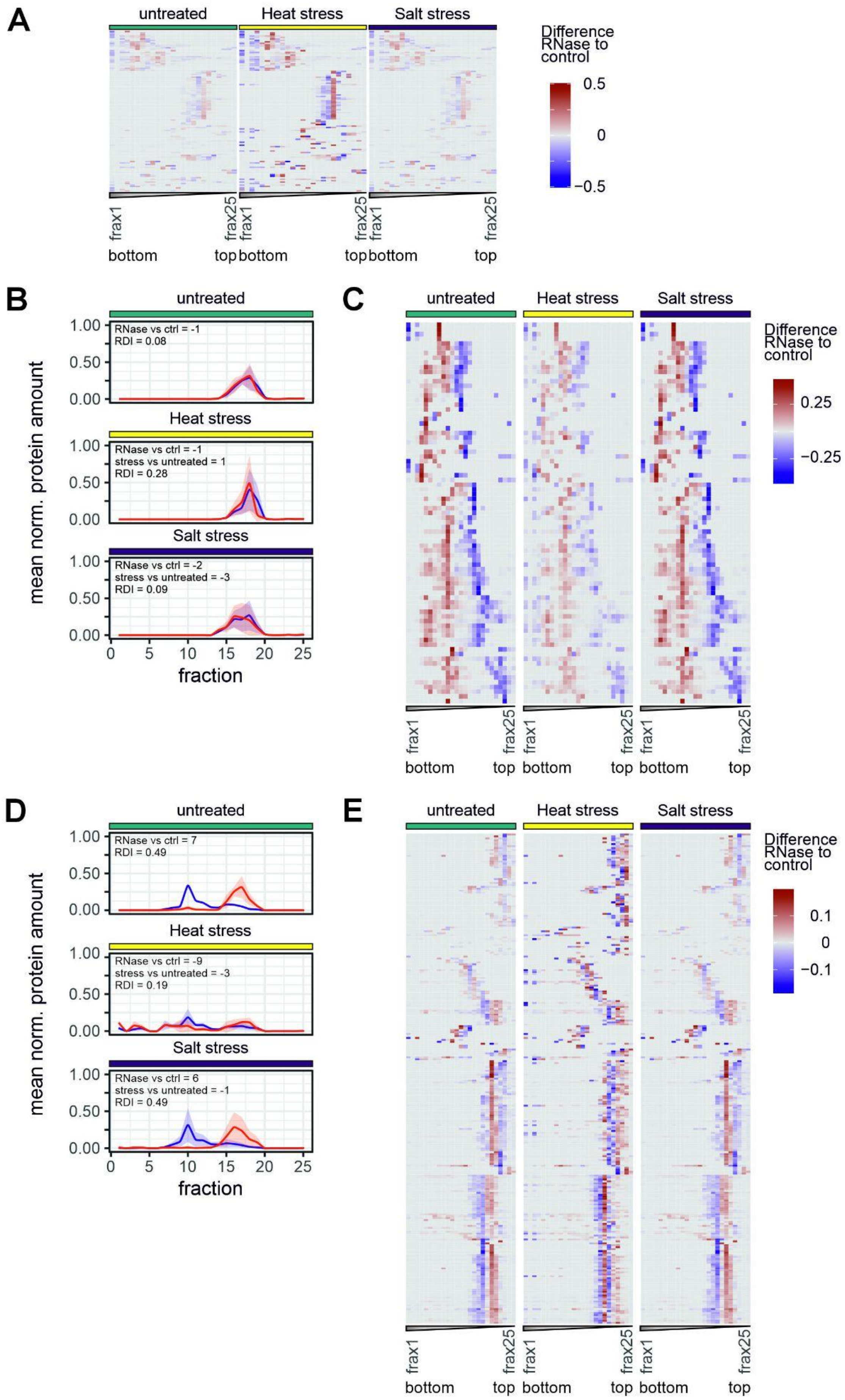
A
untreated
Heat stress
Salt stress
Difference RNase to control
0.5
0
−0.5
frax1
frax25
bottom
top
B
untreated
RNase vs ctrl = -1
RDI = 0.08
Heat stress
RNase vs ctrl = -1
stress vs untreated = 1
RDI = 0.28
Salt stress
RNase vs ctrl = -2
stress vs untreated = -3
RDI = 0.09
mean norm. protein amount
fraction
C
0.25
−0.25
D
RNase vs ctrl = 7
RDI = 0.49
RNase vs ctrl = -9
stress vs untreated = -3
RDI = 0.19
RNase vs ctrl = 6
stress vs untreated = -1
RDI = 0.49
E
0.1
0
−0.1

**Figure S3: A** Sedimentation changes for proteins in cluster 4 across all three conditions. Heatmaps depict the difference in mean normalized protein abundance between RNase-treated and control gradients across all 25 fractions (frax). Signals lost and gained after RNase treatment are indicated in red and blue, respectively. **B** Sedimentation profiles of Dys1 (P38791), a representative from cluster 4, across all three conditions. Control and RNase-treated gradients are depicted in blue and red, respectively, with shaded areas corresponding to ±1 standard deviation. Calculated RDIs and sedimentation offset values are indicated for each condition. **C** Sedimentation changes for proteins in cluster 9, as described in A. **D** Sedimentation profiles of Cam1 (P29547), a representative of cluster 9, as described in B. **E** Sedimentation changes for proteins in cluster 3, as described in A.

### Figure S4

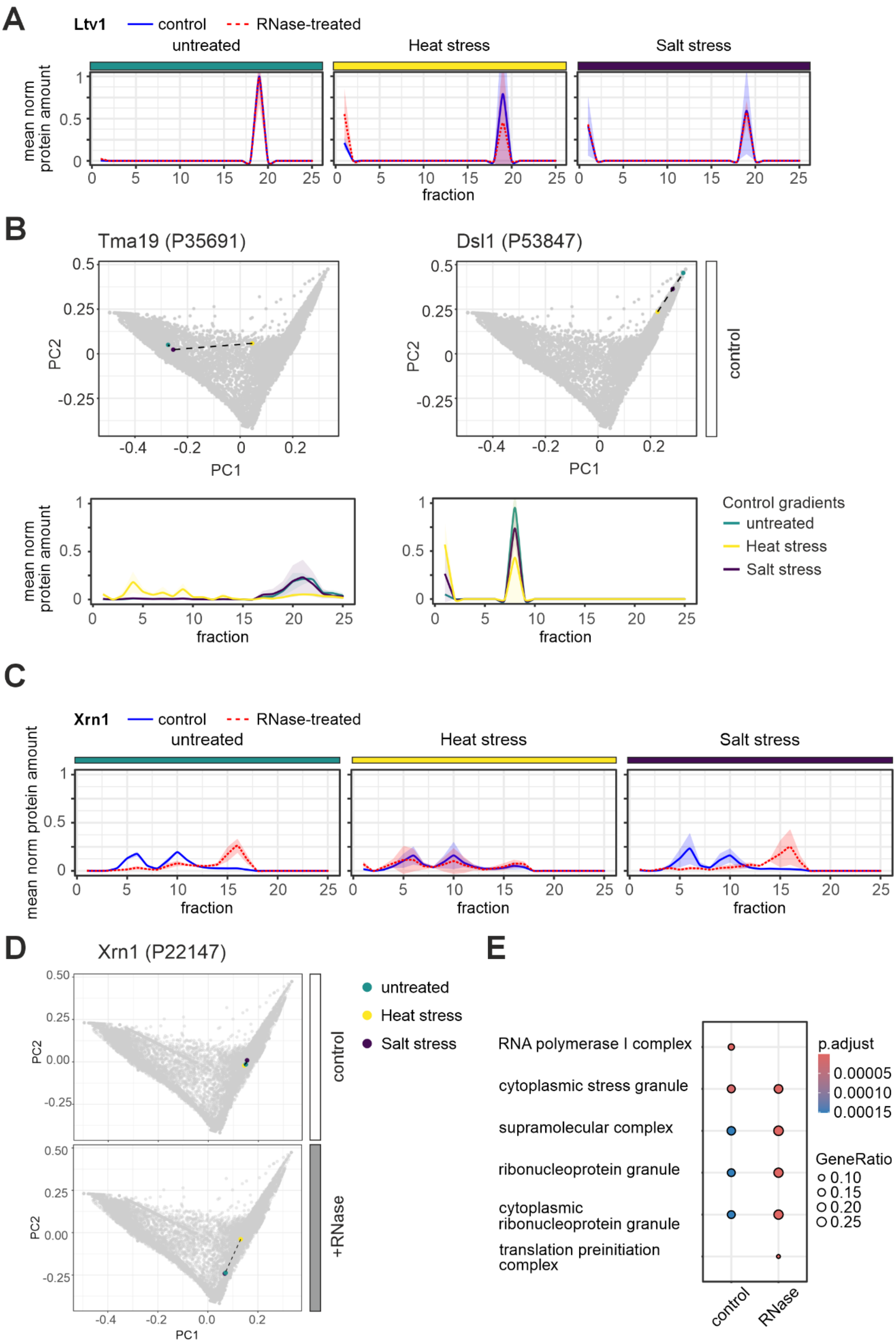
A
Ltv1
control
RNase-treated
untreated
Heat stress
Salt stress
mean norm protein amount
fraction
B
Tma19 (P35691)
Dsl1 (P53847)
PC1
PC2
control
mean norm protein amount
fraction
Control gradients
untreated
Heat stress
Salt stress
C
Xrn1
control
RNase-treated
untreated
Heat stress
Salt stress
mean norm protein amount
fraction
D
Xrn1 (P22147)
untreated
Heat stress
Salt stress
control
+RNase
PC1
PC2
E
RNA polymerase I complex
cytoplasmic stress granule
supramolecular complex
ribonucleoprotein granule
cytoplasmic ribonucleoprotein granule
translation preinitiation complex
control
RNase
p.adjust
0.00005
0.00010
0.00015
GeneRatio
0.10
0.15
0.20
0.25

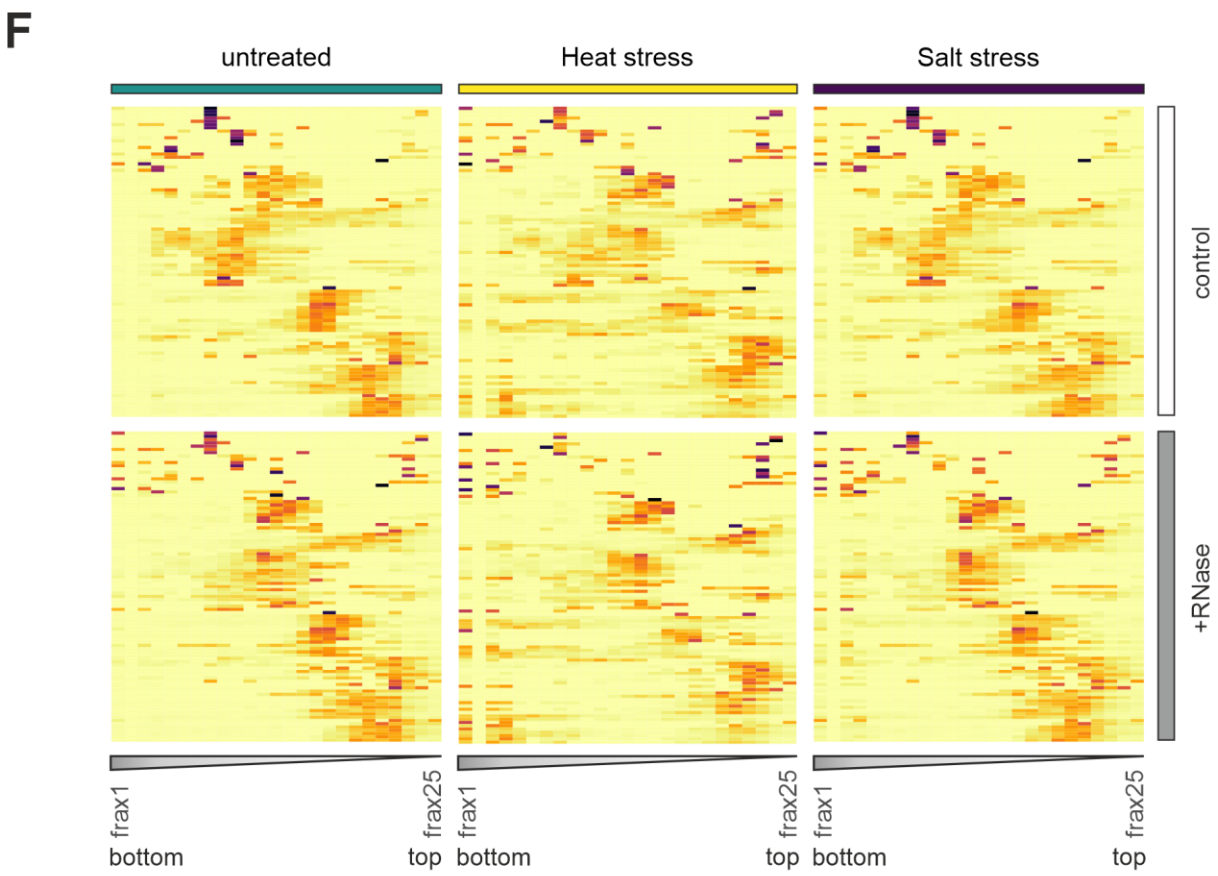


**Figure S4:** **A** Sedimentation profiles of Ltv1 across all three conditions. Control and RNase-treated gradients are depicted in blue and red, respectively, with shaded areas corresponding to ±1 standard deviation. **B** Protein-level PCA comparing individual sedimentation profiles across all three conditions (top panels), with Tma19 and Dsl1 highlighted as examples of proteins whose sedimentation in control gradients is altered after stress treatment. The corresponding sedimentation profiles across all three conditions are shown on the bottom, with untreated, heat and salt stress conditions depicted in teal, yellow and purple, respectively, with shaded areas corresponding to ±1 standard deviation. **C** Sedimentation profiles of Xrn1 across all three conditions as in A. **D** Protein-level PCA as in B, with Xrn1 highlighted. **E** GO term enrichment analysis (clusterProfiler) of proteins exhibiting altered sedimentation following heat stress, as determined by Euclidean distance. Enrichment is determined relative to the random distribution of terms across the full dataset, with node colour representing the statistical significance (p-values adjusted for multiple testing using the Benjamini-Hochberg procedure) and circle size representing term enrichment (GeneRatio), respectively. The top five cellular component terms per cluster are shown. **F** Sedimentation profiles of all proteins exhibiting heat stress-specific behaviour, defined as having a Euclidean distance greater than 0.1 between the heat-stressed gradient and the other conditions. The distribution of these heat-specific proteins is shown across all three conditions for both the control (upper panel) and RNase-treated (lower panel) gradients.

### Figure S5

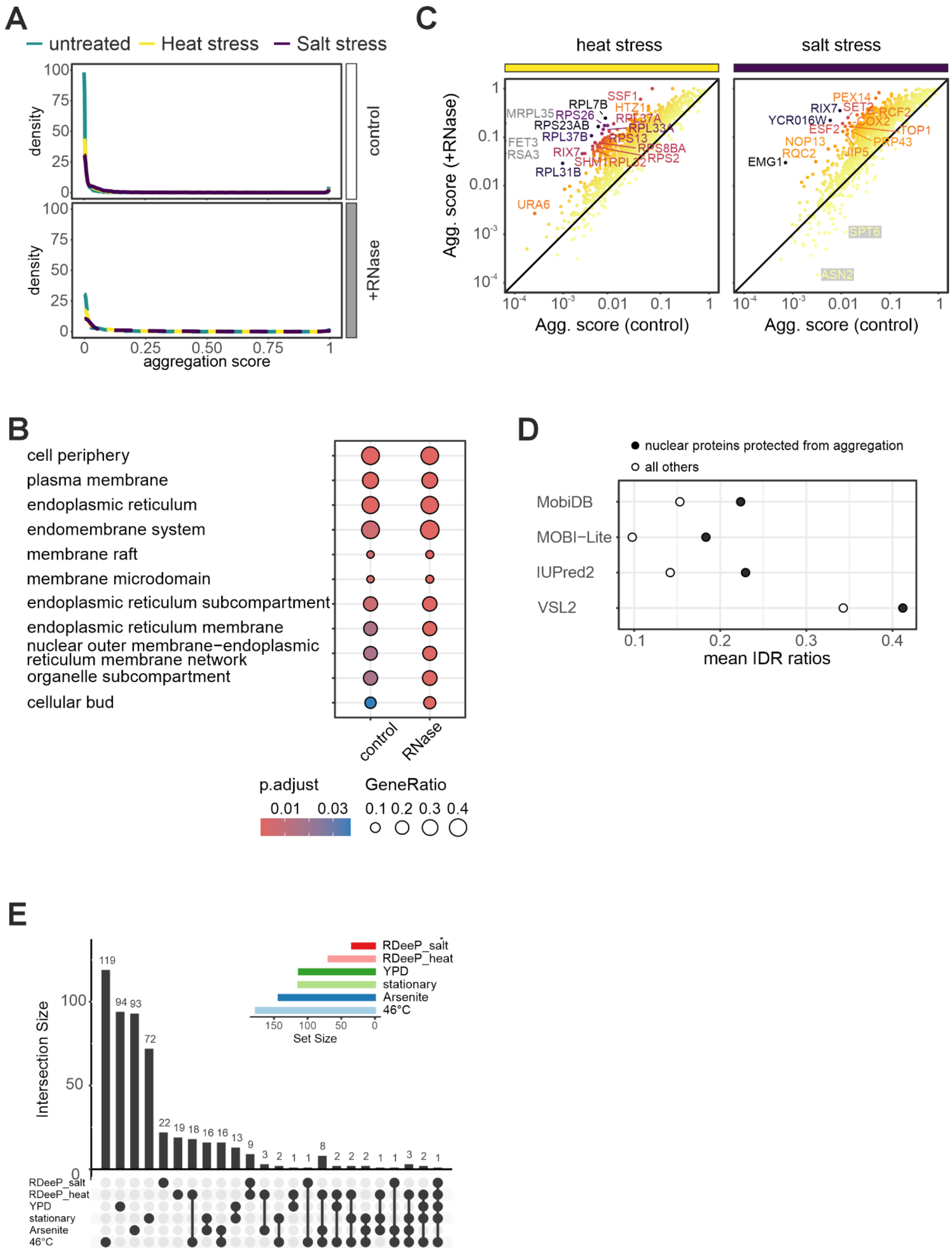


**Figure S5: A** Density distribution of the calculated aggregation scores for all detected proteins across the three conditions (untreated, heat stress, and salt stress) in both the control (upper panel) and RNase-treated (lower panel) gradients. Most proteins maintain a low basal aggregation score. **B** Gene ontology (GO) cellular component enrichment analysis for constitutively aggregating proteins, defined as those with an aggregation score > 0.5 across all three conditions in both control and RNase-treated gradients. Node color indicates statistical significance (p-values adjusted for multiple testing using the Benjamini-Hochberg procedure), and circle size represents the gene ratio. **C** Comparison of protein aggregation scores following RNase treatment of heat-stressed (left) and salt-stressed cells (right) relative to the control. Colouring represents foldchange of aggregation score after RNase treatment relative to control. **D** Comparison of median IDR ratios of nuclear proteins shielded from aggregation compared to the remaining proteins detected in the dataset; values represent consensus scores generated using MobiDB, MOBI-Lite, IUPred2, and VSL2, designed to predict and score intrinsic disorder in protein structures, which were retrieved from French-Pacheco et al., 2022. **E** Upset plot detailing the overlap of the stress-specific aggregators (heat or salt) identified in this study with previously published yeast aggregation datasets, including nutrient limitation (stationary phase; Narayanaswamy et al., 2009), arsenite stress (Ibstedt et al., 2014), and severe 46°C heat stress (Wallace et al. 2015). Baseline aggregation in rich medium (YPD) was taken from Ibstadt et al., 2014.

### Figure S6

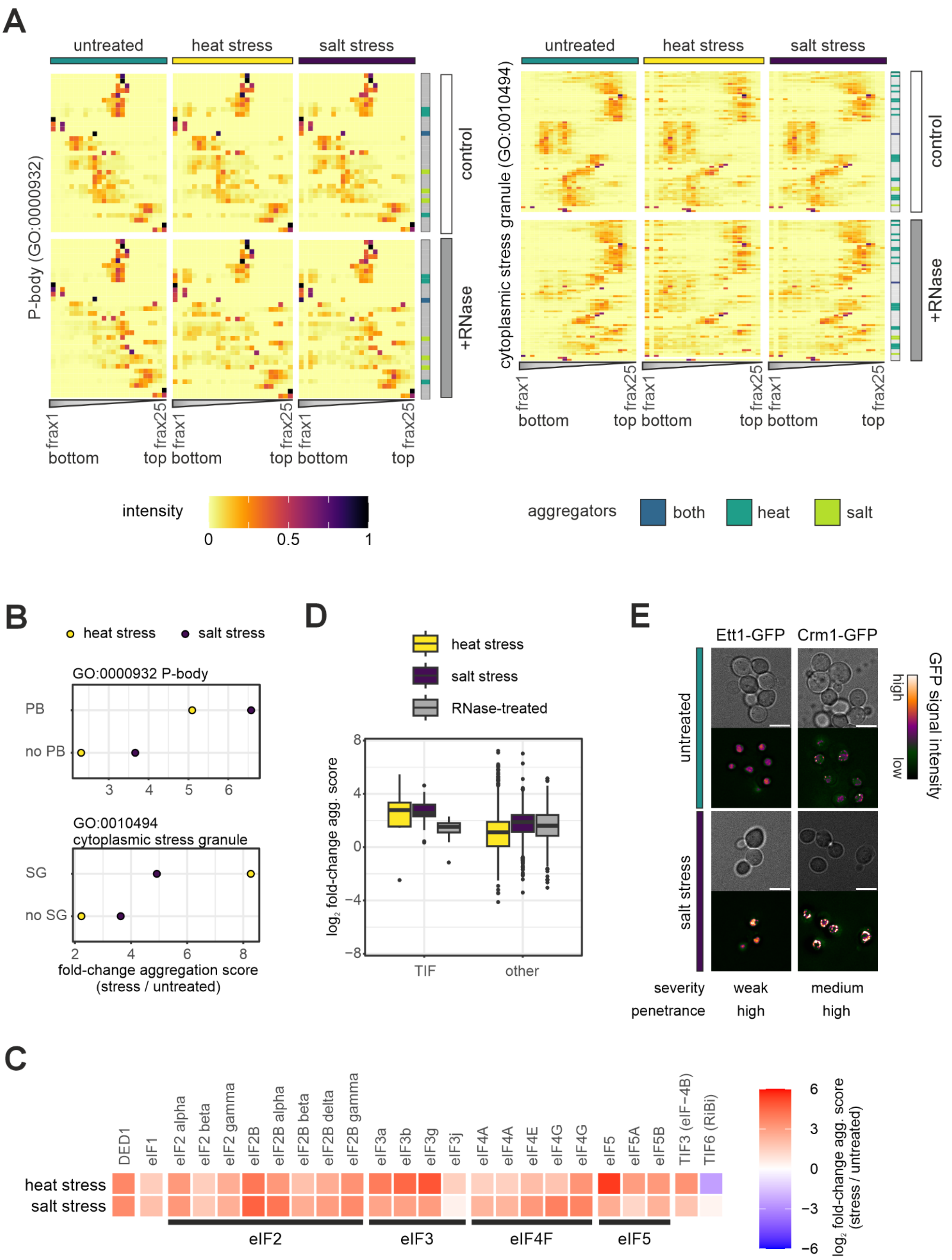


**Figure S6: A** Distribution of P-body (upper panels) and stress granule components (lower panels) in control and RNase-treated gradients across all three conditions. Stress-specific aggregators are highlighted to the right in dark or light green and blue for aggregation in response to heat, salt, or under both conditions, respectively. **B** Median foldchange of aggregation score for proteins associated with the GO terms P-body (GO:0000932) (upper panel) or cytoplasmic stress granule (GO:0010494) (lower panel) in comparison to the remaining proteins in the dataset. **C** Foldchange increase in aggregation scores after heat and salt stress for proteins with translation initiation factor activity (GO:0003743). TIF6 has similarity to human translation initiation factor 6 (eIF6) but is involved in ribosome biogenesis (RiBi) in yeast. **D** Comparison of log2 fold-change aggregation scores for proteins with translation initiation factor activity (GO:0003743) and all other proteins with non-zero aggregation scores for heat stress (yellow), salt stress (purple), and RNase treatment (grey), all relative to the untreated control. **E** Representative live-cell fluorescence microscopy images of the GFP-tagged Ett1 and Crm1, comparing untreated cells with cells subjected to salt stress (n = 3 biological replicates). The relative severity and penetrance of the observed condensation phenotypes are summarized below the images. GFP signal intensity is visualized using a false-colour scale (Blue Orange icb), as indicated to the right. Contrast and brightness settings were kept constant between untreated and salt-stressed conditions for each protein. Scale bars correspond to 5 µm.
